# CDC5L surveils cellular stress responses and stress granule formation through transcriptional repression

**DOI:** 10.1101/2024.01.04.574133

**Authors:** Beituo Qian, Shunyi Li, Yongjia Duan, Feng Qiu, Rirong Hu, Wenkai Yue, Jihong Cui, Qiangqiang Wang, Wanjin Li, Yanshan Fang

## Abstract

Cells have evolved a variety of mechanisms to respond to stress, such as activating the PERK– eIF2α pathway and forming stress granules (SGs). It is important that these mechanisms are inducted only when necessary and exerted at appropriate levels, to prevent spontaneous or excessive activation of stress responses. However, the mechanisms by which cells keep the stress response programs in check are elusive. In this study, we discovered that downregulation of *Cell Division Cycle 5 Like* (*CDC5L*) causes spontaneous SG formation in the absence of any stress, which is independent of its known functions in the cell cycle or the PRP19 complex. Instead, we found that CDC5L binds to the *PERK* promoter through its DNA-binding domains and represses *PERK* mRNA transcription. As a result, it negatively regulates the abundance of PERK protein and the phosphorylation levels of eIF2α, thereby suppressing the PERK–eIF2α signaling pathway and preventing undesirable SG assembly. Further RNA-sequencing (seq) and chromatin immunoprecipitation (ChIP)-seq analyses reveal a dual function of CDC5L in gene transcription: it acts as a transcriptional activator in cell cycle control but as a repressor in cellular stress responses. Finally, we show that the loss of *CDC5L* decreases cell viability and fly survival under mild stress conditions. Together, our findings demonstrate a previously unknown role and mechanism of CDC5L in the surveillance of cellular stress through transcriptional repression, which serves as a gatekeeper for the stress response programs such as the PERK–eIF2α pathway and SG formation.

**Significance statement:** Cells need to respond to stress promptly for survival. Meanwhile, it is equally important to prevent spontaneous or excessive activation of stress response programs when no stress or only minor stress is present. Here, we reveal that the DNA/RNA-binding protein CDC5L represses the transcription of a cluster of stress response genes including *PERK*. In doing so, CDC5L suppresses the PERK-eIF2α pathway and prevents spontaneous SG assembly. Downregulation of *CDC5L* releases the restraint on these genes, resulting in an exaggerated response to stress and decreased viability in both cell and fly models. Taken together, this study demonstrates the existence of a gatekeeper mechanism that surveils the stress response programs and highlights the crucial role of CDC5L-mediated transcriptional repression in this regulation.

## INTRODUCTION

Diverse cellular stress can occur throughout an individual’s lifetime. The ultimate fate of a stressed cell depends on the type and severity of the stress as well as the cell’s ability to promptly and properly cope with it. For instance, oxidative stress, heat shock, and other stressors can lead to rapid assembly of SGs, which are then dismantled when the stress is relieved. The dynamic assembly and diassembly of SGs are believed to promote cell survival during stress and are implicated in various physiological and pathophysiological processes (Alberti and Hyman, 2021; Glauninger et al., 2022).

SGs are membraneless biomolecular condensates that contain RNA and phase-separated proteins, particularly RNA-binding proteins (RBPs) (Protter and Parker, 2016; Alberti and Hyman, 2021; Roden and Gladfelter, 2021; Glauninger et al., 2022). For example, the Ras-GTPase-activating protein SH3 domain-binding protein (G3BP) serves as a central RBP of SGs, and depleting it abolishes SG formation (Guillen-Boixet et al., 2020; Yang et al., 2020; Gwon et al., 2021). Additionally, various nuclear RBPs such as T-cell intracellular antigen 1-related (TIAR), TAR DNA-binding protein 43 (TDP-43) and fused in sarcoma (FUS) are recruited from the nucleus to the cytoplasm, participating in SG assembly (Ravanidis et al., 2018; Youn et al., 2019; Portz et al., 2021). These SG-associated RBPs are often detected in pathological protein inclusions present in patients with amyotrophic lateral sclerosis (ALS) or frontotemporal dementia (FTD). Furthermore, disease-causing mutations have been identified in the genes encoding these proteins. As a result, the aberrant formation and transition of liquid droplet-like SGs to solid protein aggregation are believed to play a significant role in the pathogenesis of these diseases (Wolozin and Ivanov, 2019; Baradaran-Heravi et al., 2020; Alberti and Hyman, 2021). Hence, it is crucial to prevent undesired SG assembly in the first place.

A well-known signaling pathway that triggers SG formation is the protein kinase RNA-like endoplasmic reticulum kinase (PERK)–eukaryotic initiation factor 2 alpha (eIF2α) pathway. A variety of cellular stressors, such as endoplasmic reticulum (ER) stress, heat-shock stress, and arsenite-induced oxidative stress, can activate PERK, a kinase that phosphorylates eIF2α. Consequently, the phosphorylation levels of eIF2α increase rapidly, leading to the inhibition of mRNA translation and the initiation of SG assembly. The translation initiation factors, 40S ribosomal subunits, untranslated mRNAs and some RBPs condense to form the core of SGs, which further incorporates additional RBPs to become mature SGs (Anderson and Kedersha, 2009; Buchan and Parker, 2009). And, it is shown that modulation of eIF2α phosphorylation levels, for example, by using PERK inhibitors, can regulate the assembly-disassembly dynamics of SGs (Zhang et al., 2018b; Fang et al., 2019; Hans et al., 2020).

Various chemicals, such as oxidative stress-inducing arsenite and proteasome inhibitor bortezomib, as well as harsh physical conditions like heat-shock stress, are commonly employed in laboratories to induce SGs. In addition, overexpression (OE) of G3BP1 is sufficient to trigger SG assembly (Guillen-Boixet et al., 2020; Yang et al., 2020). However, it is worth noting that SGs are seldom associated with loss-of-function (LOF) mutations in genes, which raises doubts regarding whether the formation of SGs is an intrinsic, genetically-programed cellular function (Alberti et al., 2019; Glauninger et al., 2022; Putnam et al., 2023). Here, we report that downregulation of *CDC5L*, a highly conserved gene in eukaryotes, leads to spontaneous SG formation. *CDC5L* encodes a DNA/RNA-binding protein that was initially identified as an essential gene for G2/M progression in the cell cycle (Bernstein and Coughlin, 1998). Subsequently, it was found as a major component of the pre-mRNA-processing factor 19 (PRP19) complex, which is involved in various cellular processes such as splicing, transcription, and mRNA export (Ajuh et al., 2000; Boudrez et al., 2000; Chanarat and Strasser, 2013). Surprisingly, we find that the role of CDC5L in regulating SGs is independent of its known functions in the cell cycle or the PRP19 complex. Instead, CDC5L functions as a transcriptional repressor, negatively regulating the PERK-eIF2α pathway to suppress the undesired activation of stress responses and SG assembly.

## RESULTS

### Downregulation of *CDC5L* causes spontaneous SG formation and enhances SG assembly in response to stress

A recent bioinformatic analysis of protein-protein interaction networks identified *CDC5L* as one of the major hub genes of ALS-related proteins, many of which are localized to SGs and/or associated with stress responses or cellular homeostasis (Kumar and Haider, 2022). However, the specific connection between CDC5L and SGs, as well as the underlying mechanism, remains unknown. In this study, our objective was to investigate the involvement of CDC5L in the regulation of SGs or cellular stress responses.

First, we knocked down *CDC5L* in HeLa cells using two independent small interfering RNAs (siRNAs) #1 and #2 (Figure S1A-B). Compared to the cells treated with the scrambled siRNA control (si-Ctrl), cells treated with the siRNAs against *CDC5L* exhibited increased incidence of spontaneous formation of G3BP1+ granules in the cytoplasm (Figure 1A-B). Since the two siRNAs targeting *CDC5L* showed similar knockdown (KD) efficiencies and induction effects on SG formation, we used the siRNA-#2 for the rest of the study and referred to it as si-*CDC5L*. We further confirmed that si-*CDC5L* induced the formation of SGs by immunostaining with another commonly used SG marker, TIAR (Figure S1C).

**Figure 1.**
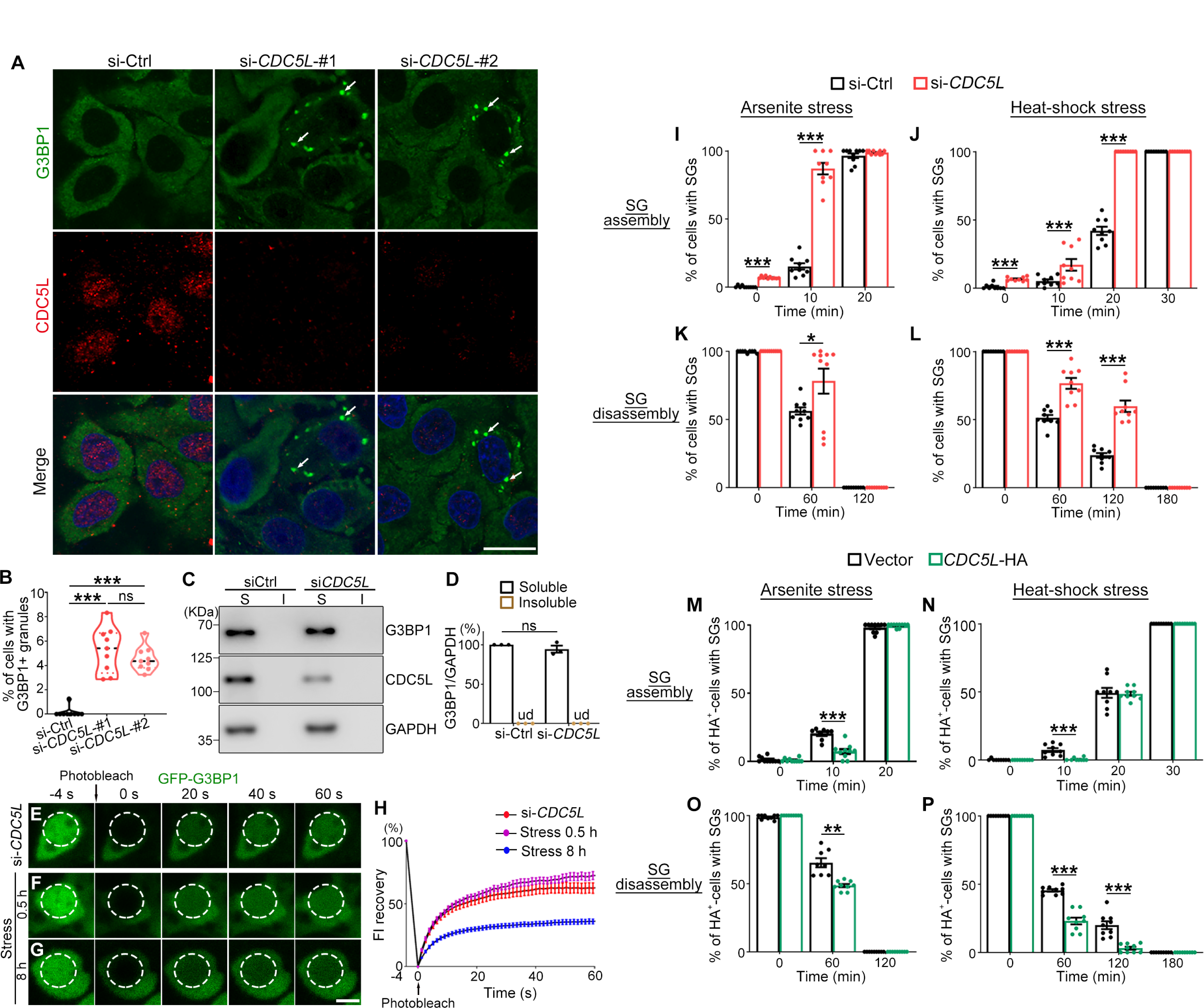
KD of *CDC5L* induces spontaneous SG formation and accelerates stress-induced SG assembly. (**A**-**B**) Representative confocal images (A) and quantification (B) of HeLa cells transfected with scrambled siRNA as a control (si-Ctrl) or two independent siRNAs against *CDC5L* (si-*CDC5L*-#1 and -#2). All cells are stained for G3BP1, CDC5L and DAPI (for nucleus). Arrows, G3BP1+ granules. (**C-D**) Western blot analysis (C) and quantification (D) of the soluble (S, supernatants in RIPA) and insoluble (I, pellets resuspended in 9 M urea) fractions of the lysates of HeLa cells transfected with indicated siRNAs. All protein levels are normalized to GAPDH in the soluble fraction. (**E**-**H**) Representative images (E-G) and quantification (H) of the FRAP analysis of the fluorescence intensity (FI) of GFP-G3BP1 granules induced by KD of *CDC5L* (E), acute cellular stress (250 μM arsenite, 0.5 h) (F), or prolonged cellular stress (250 μM arsenite, 8 h) (G) in *live* cells. (**I-P**) The impact of KD (I-L) or OE (M-P) of *CDC5L* on the kinetics of SG assembly and disassembly is assessed by quantification of the average percentages of cells with SGs induced by arsenite stress (250 μM) (I, K, M, O) or heat shock (42°C) (J, L, N, P) at the indicated time points after stress or during recovery (after arsenite washout or returning to 37°C; see Figure S2 and S3 for the representative images). Data are shown in violin plots (B) or as mean ± SEM (D, H-P). n ≥ 200 cells in each group from pooled results of 3 independent repeats (B, I-P), n = 3 (D), and n ≥ 18 puncta in each group in (H). The statistical significance is determined by one-way ANOVA (B) or Student’s *t*-test (D, I-P) with \**p* < 0.05, \*\**p* < 0.01, \*\*\**p* < 0.001; ns, not significant. ud, undetected. Scale bars: 20 μm in (A) and 2 μm in (E-G).

Next, we examined and showed that KD of *CDC5L* did not change the solubility of G3BP1 protein (Figure 1C-D), which excluded the possibility that the si-*CDC5L*-induced cytoplasmic puncta were protein aggregates containing G3BP1. Further, we conducted the fluorescence recovery after photobleaching (FRAP) analysis in *live* cells using green fluorescent protein (GFP)-tagged G3BP1. The fluorescence intensity (FI) of the si-*CDC5L*-induced GFP-G3BP1 granules rapidly recovered after photobleaching (Figure 1E, 1H), and the kinetics was similar to that induced by acute arsenite stress (250 μM, 0.5 h) (Figure 1F, 1H), suggesting that the GFP-G3BP1 granules in both instances were dynamic and liquid-like. In contrast, under prolonged stress (250 μM, 8 h), SGs were solidified, as evidenced by the markedly reduced FI recovery (Figure 1G-H). Together, these data indicate that loss of *CDC5L* causes spontaneous SG formation in cells.

Moreover, we found that the levels of *CDC5L* also modulated the assembly-disassembly dynamics of SGs in response to stress. Our data showed that for both arsenite stress (250 μM; Figure S2) and heat-shock stress (42°C; Figure S3), KD of *CDC5L* promoted SG assembly and hindered SG disassembly (Figure 1I-L), while OE of *CDC5L* delayed SG assembly and accelerated SG disassembly (Figure 1M-P).

### The impact of *CDC5L* on SGs is independent of the cell cycle or the PRP19 complex

To understand how *CDC5L* regulates SG formation, we first examined how the CDC5L protein responded to stress. Unlike the SG marker TIAR, which was recruited from the nucleus to SGs in the cytoplasm upon arsenite stress, CDC5L was predominantly nuclear under normal conditions and remained in the nucleus during stress (Figure S4A). We then showed that the stress signaling pathway was indeed activated by arsenite stress, evidenced by increased phosphorylation levels of eIF2α (Figure S4B-S4D). In contrast, the protein levels of CDC5L was unaffected (Figure S4B and S4E). Given that increase of G3BP1 levels could induce SG assembly (Guillen-Boixet et al., 2020; Yang et al., 2020), we examined the protein levels of G3BP1 in cells with si-*CDC5L* and no significant change was detected (Figure 1C-D), which excluded the possibility that KD of *CDC5L* promoted SG assembly by increasing G3BP1 levels.

CDC5L was known to play an important role in the cell cycle (Bernstein and Coughlin, 1998; Williams et al., 2006). Since KD of *CDC5L* might perturb the natural cell cycle and hinder G2/M progression, we wondered if spontaneous SG formation was associated with any specific phase in the cell cycle, particularly the G2/M phase. We then synchronized the phase of HeLa cells (Figure 2A; also see Methods and Figure S5) and scrutinized these cells for SGs by immunostaining with an anti-G3BP1 antibody. However, no spontaneous SG was found in the G1, S or G2/M phase (Figure 2B-C). Furthermore, we showed that KD of *CDC5L* could cause spontaneous SG formation in all the above phases, and the incidence rate was similar in the different phases (Figure 2D-E). Of note, no SG was detected in cells that underwent active mitosis, which is consistent with the previous report that cells in the metaphase of mitosis did not form SGs even in the presence of arsenite stress (Sivan et al., 2007). Thus, an increase in the proportion of cells in the G2/M phase, for instance, by loss of *CDC5L* (Bernstein and Coughlin, 1998), would decrease, but not increase, the likelihood of SG formation, which ruled out the possibility that the function of *CDC5L* in the cell cycle accounts for the si-*CDC5L*-induced spontaneous formation of SGs.

**Figure 2.**
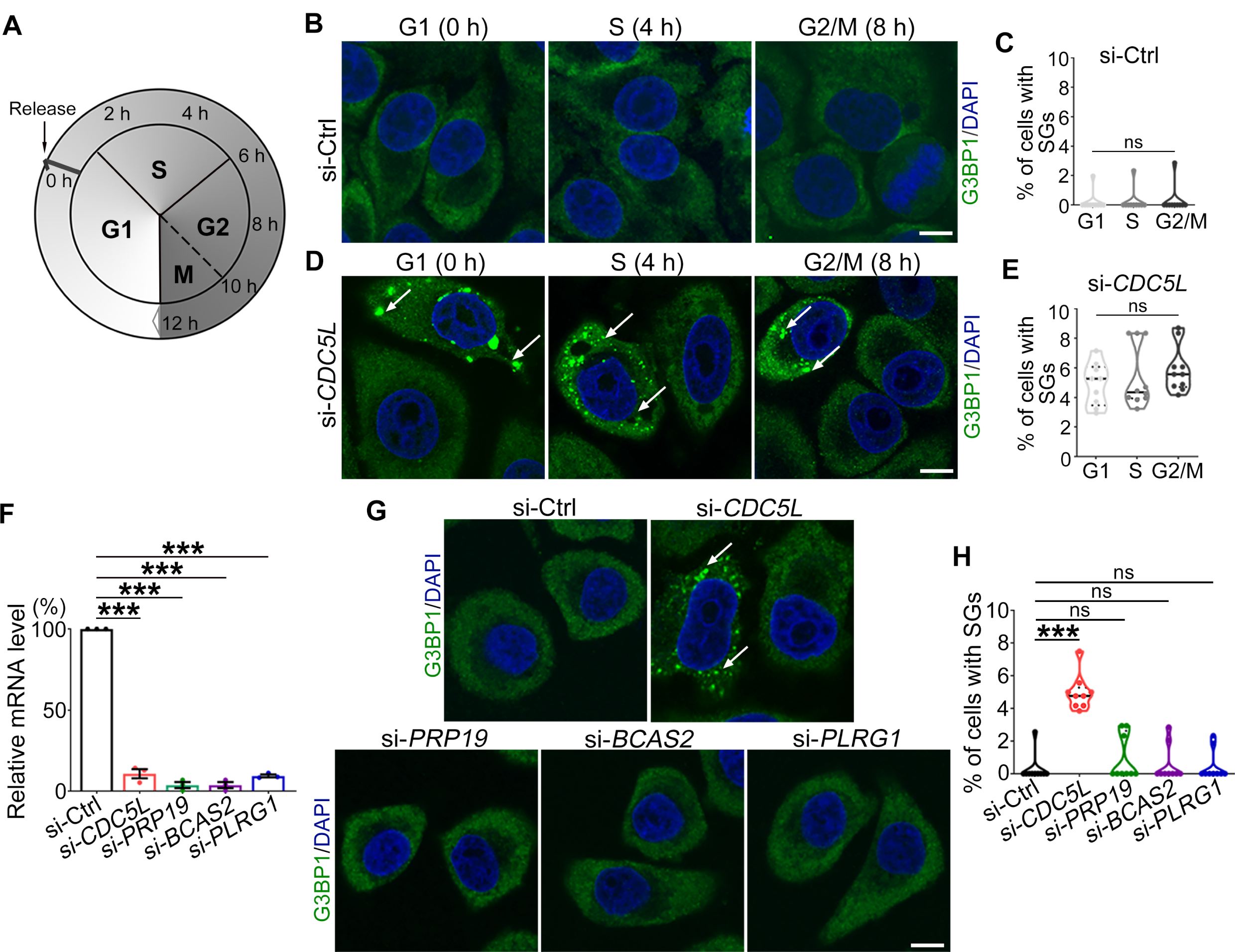
No spontaneous SG assembly during the cell cycle or by KD of other main components of the PRP19 complex. (**A**) A diagraph of the cell cycle assay. HeLa cells are synchronized to the G1 phase and then released. The time and the corresponding phases of the cell cycle are indicated (see Methods and Figure S5). (**B**-**C**) No spontaneous SG assembly is observed in any phase of the cell cycle. (**D-E**) si-*CDC5L*-induced spontaneous SGs are found in all phases of the cell cycle at similar occurrence rates. G3BP1, SGs (arrows); DAPI, nucleus. (**F**) qPCR analysis confirming the KD efficiency of the siRNAs against the indicated main components of the PRP19 complex. All mRNA levels are normalized to *GAPDH* and shown as the average percentage to that of the control group (si-Ctrl, scrambled siRNA). (**G**-**H**) Representative confocal images (G) and quantification (H) of the average percentage of cells forming spontaneous SGs with the indicated siRNAs. Data are shown in violin plots (C, E, H) or as mean ± SEM (F). n ≥ 200 cells in each group from pooled results of 3 independent repeats (C, E, H) and n = 3 (F). One-way ANOVA; \*\*\**p* < 0.001; ns, not significant. Scale bars: 10 μm.

Another well-known function of CDC5L is its participation in the PRP19 complex, which regulates pre-mRNA splicing, transcription elongation, DNA repair, and other cellular processes (Boudrez et al., 2000; Chanarat and Strasser, 2013). We then knocked down each main component of the PRP19 complex, including PRP19, BCAS2 and PLRG1, in addition to CDC5L (Figure 2F). However, none of the remaining proteins manifested the same effect as KD of *CDC5L* to induce spontaneous SG formation (Figure 2G-H). Together, these results indicate that the role of CDC5L in regulating SGs is rather unique and not mediated by the PRP19 complex, suggesting an unexplored function and molecular mechanism of CDC5L.

### CDC5L regulates PERK protein abundance and the basal phosphorylation levels of eIF2α

Next, we asked whether CDC5L affected the stress reponse pathways, such as the PERK–eIF2α signaling pathway, which is activated in resposne to arsenite or heat-shock stress and initiates SG assembly (Anderson and Kedersha, 2009; Buchan and Parker, 2009). Interestingly, we found that both the protein abundance of PERK and the phosphorylaiton levels of eIF2α were significantly increased by *CDC5L* KD (Figure 3A-C), indicating an elevated basal level of the PERK–eIF2α pathway. We confirmed that increase of PERK protein levels by transient OE of *PERK*-HA could increase the phosphorylation levels of eIF2α (Figure 3D-E) and was sufficient to induce SG assembly (Figure 3F-G). Furthermore, we demonstrated that KD of *PERK* substantially reduced the increase of eIF2α phosphorylation caused by *CDC5L* KD (Figure 3H-J) and abolished the spontaneous SG formation with si-*CDC5L* (Figure 3K-L). Together, these results suggest that CDC5L modulates the basal levels of the PERK-eIF2α pathway, and its abnormal upregulation underlies the spontaneous SG formation in cells with loss of *CDC5L*.

**Figure 3.**
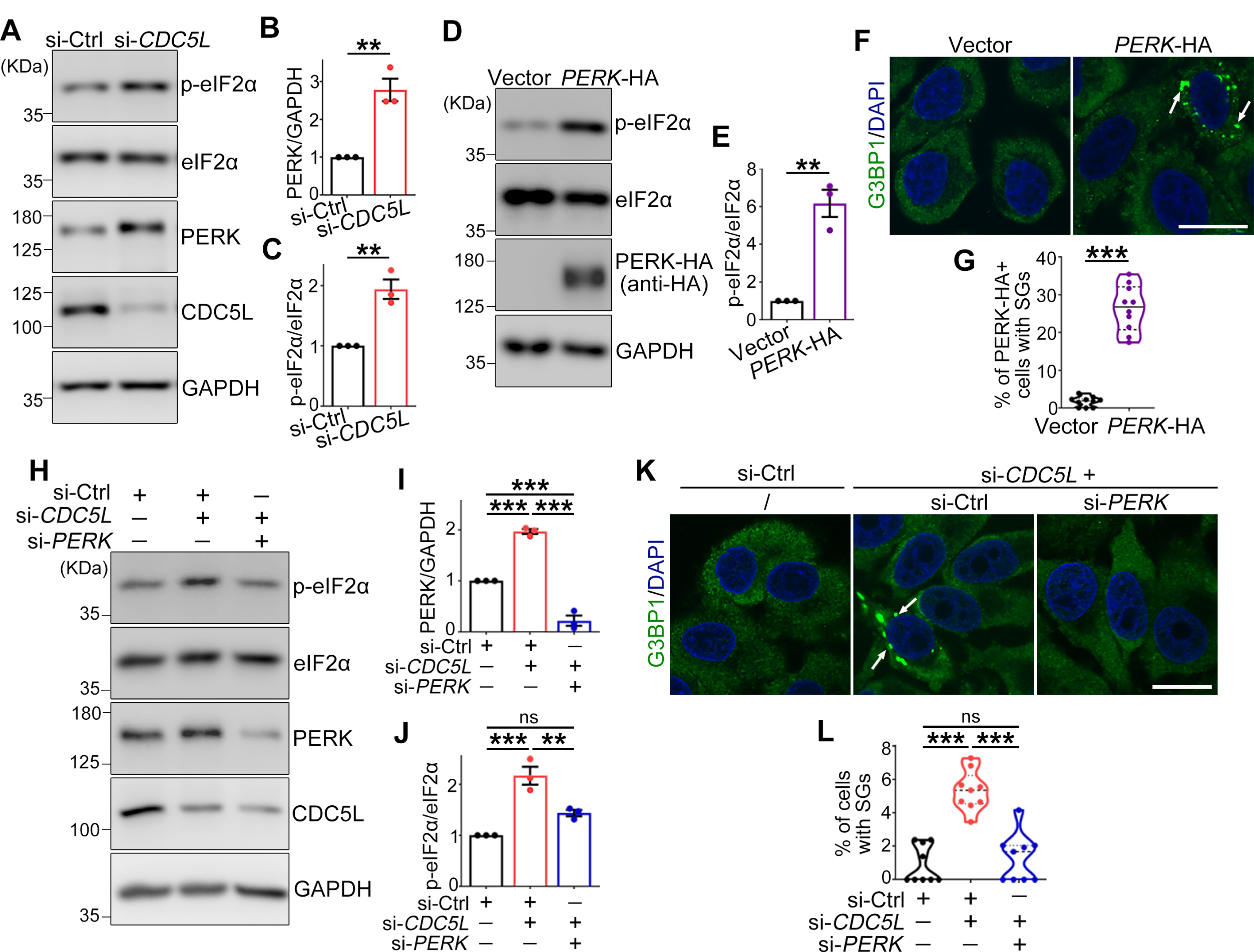
The level of the PERK–eIF2α pathway is markedly elevated and this increase underlies the *CDC5L* KD-induced SG formation. (**A-E**) Representative images of the western blots (A, D) and quantifications of PERK protein levels (B) and eIF2α phosphorylation levels (C, E) in HeLa cells with *CDC5L* KD (A-C) or *PERK* OE (D-E). p-eIF2α, phosphorylated eIF2α. (**F-G**) Representative confocal images (F) and quantification (G) of the percentage of cells forming SGs with *PERK* OE. G3BP1, SGs (arrows); DAPI, nucleus. (**H-J**) Western blot analysis (H) and quantifications of PERK protein levels (I) and eIF2α phosphorylation levels (J) in HeLa cells with *CDC5L* KD in the absence or presence of *PERK* KD. (**K-L**) Representative confocal images (K) and quantification (L) of the percentage of cells forming SGs with *CDC5L* KD in the absence or presence of *PERK* KD. Mean ± SEM (B-C, E, I-J) or violin plots (G, L). n = 3 in (B-C, E, I-J) and n ≥ 200 cells in each group from pooled results of 3 independent repeats in (G, L). Student’s *t*-test in (B-C, E, G) and one-way ANOVA in (I-J, L); \*\**p* < 0.01, \*\*\**p* < 0.001; ns, not significant. Scale bars: 20 μm.

### CDC5L binds to the *PERK* promoter and represses *PERK* mRNA transcription

To understand why PERK protein abundance was affected by *CDC5L* KD, we examined the mRNA levels of *PERK* using quantitative real-time PCR (qPCR). The results showed that *CDC5L* KD dramatically increased the mRNA levels of *PERK* (Figure 4A). In contrast, the mRNA levels of three other kinases known to phosphorylate eIF2α—protein kinase double-stranded RNA-dependent (PKR), general control non-derepressible-2 (GCN2), and heme-regulated inhibitor (HRI) (Donnelly et al., 2013)—were not significantly changed by *CDC5L* KD (Figure S6). This data indicates that CDC5L specifically affected *PERK* among the four eIF2α kinases. Consistently, *CDC5L* OE decreased the mRNA levels of *PERK*, although to a moderate extent (Figure 4B), which suggests that *PERK* expression was constantly and adequately repressed in cells under normal conditions.

**Figure 4.**
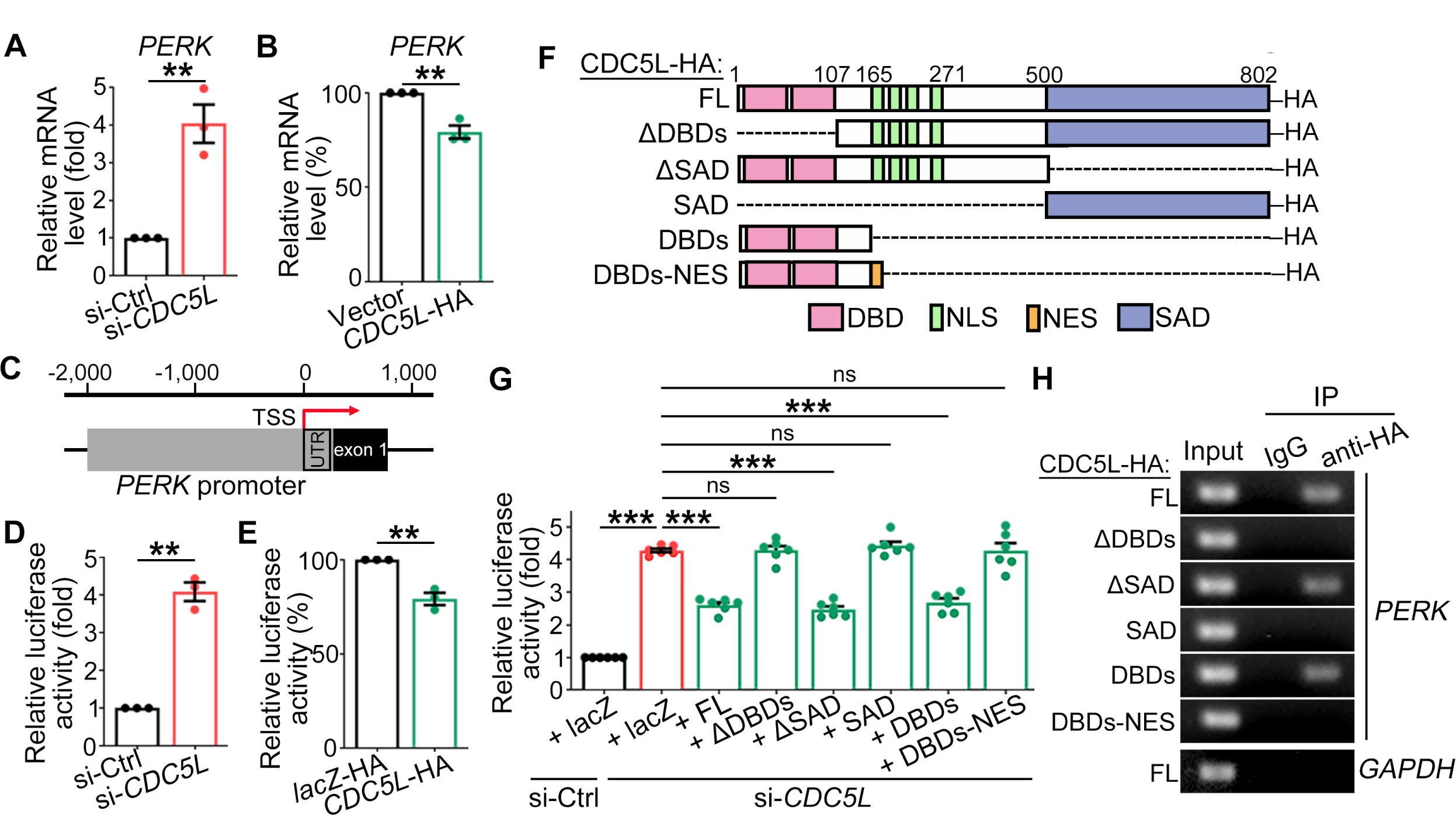
CDC5L represses *PERK* mRNA transcription through its DBDs. (**A-B**) The qPCR analysis of the mRNA levels of *PERK* in HeLa cells with *CDC5L* KD (A) or OE (B). The mRNA levels are normalized to *GAPDH* and shown as fold change or percentage to that of the control group (set to 1 or 100%). (**C**) A schematic diagram showing the predicted promoter region of the human *PERK* gene. Red arrow, transcription start site (TSS). (**D-E**) Relative luciferase activity of *PERK-luc* in HeLa cells with *CDC5L* KD (D) or OE (E). Scrambled siRNA (si-Ctrl) and *lacZ*-HA are used as the KD and OE control in the luciferase reporter assay, respectively (the same below). (**F**) Diagrams showing the major functional domains of the full-length (FL) CDC5L protein and the different truncated CDC5L. DBDs, DNA-binding domains; SAD, spiceosome-assoicated domain; NLS, nuclear localization sequence; NES, nuclear export signal. (**G**) Relative luciferase activity of *PERK-luc* with *CDC5L* KD in HeLa cells, co-transfected with the FL or truncated CDC5L as indicated. (**H**) ChIP for CDC5L-HA followed by semiquantitative PCR for the promoter of *PERK* or *GAPDH* (negative control) are performed in HeLa cells overexpressing the FL or truncated CDC5L-HA. Mean ± SEM; n = 3 in (A-B, D-E) and n = 6 in (G). Student’s *t*-test in (A-B, D-E) and one-way ANOVA in (G); \*\**p* < 0.01, \*\*\**p* < 0.001; ns, not significant.

To examine whether CDC5L regulated *PERK* mRNA transcription, we performed a dual luciferase assay (Sherf et al., 1996). In brief, we amplified the predicted *PERK* promoter, the DNA fragment from ∼2 kb upstream to ∼200 bp downstream of the transcription start site (TSS) of *PERK* (chr2: 88, 627,275 - 88,629,464) (Figure 4C) and fused it to the firefly luciferase reporter gene (*PERK*-*luc*), which was co-transfected into HeLa cells with a vector to express Renilla luciferase (*Rluc*) as an internal control. The data indicated that *CDC5L* KD drastically increased (Figure 4D) whereas *CDC5L* OE modestly suppressed (Figure 4E) *PERK-luc* expression, consistent with the changes in *PERK* mRNA levels by *CDC5L* KD or OE (Figure 4A-B).

The CDC5L protein contains two DNA-binding domains (DBDs) at the N-terminus, a spliceosome-associated domain (SAD) at the C-terminus, and several putative nuclear localization sequences (NLS) in the middle (Bernstein and Coughlin, 1998; van Maldegem et al., 2015). To determine what domain(s) within the CDC5L protein was required for regulating *PERK* transcription, we generated various constructs to express truncated CDC5L proteins, including the ΔDBDs (aa 107-802), ΔSAD (aa 1-500), SAD (aa 500-802), and DBDs (aa 1-165) of CDC5L-HA (Figure 4F), and compared them to the full-length (FL) CDC5L-HA using the luciferase reporter assay. Since OE of *CDC5L* exhibited only mild effects (Figure 4B and 4E), we instead examined them in a rescue experiment in the *CDC5L* KD background. si-*CDC5L* caused a robust upregulation of *PERK-luc*, which was significantly reduced by OE of FL as well as ΔSAD and DBDs, but not ΔDBDs or SAD, of the CDC5L-HA protein (Figure 4G). The result that SAD domain was dispensable for CDC5L to regulate *PERK* transcription suggested that the spliceosome-associated function of CDC5L was not required in this regulation, which was consistent with our earlier observation that KD of the other main component of the PRP19 splicing complex did not phenocopy the effect of *CDC5L* KD on SG formation (Figure 2F-H).

The DBDs of CDC5L were sufficient and necessary for the repression of *PERK* transcription (Figure 4G), which prompted us to test whether the FL and truncated CDC5L-HA proteins could bind to the *PERK* promoter. We then performed the chromatin immunoprecipitation (ChIP) and semiquantitative PCR assay. Compared to the normal rabbit IgG control, ChIP for CDC5L-HA with an anti-HA antibody (rabbit) enriched the *PERK* promoter from cells expressing FL, ΔSAD and DBDs, but not ΔDBDs or SAD, of the CDC5L-HA protein (Figure 4H). We noted that despite the change in the sub-nuclear distributions, the DBDs-HA construct (which lacked the putative NLS) was still mostly expressed in the nucleus (Figure S7). Hence, we attached a nuclear export signal (NES) (Sun et al., 2021) to the C-terminus of the DBDs to generate a DBDs-NES-HA construct. As expected, the DBDs-NES-HA was localized to the cytoplasm (Figure S7E-S7F), which abolished the repressing effect of the DBDs-HA on *PERK* transcription (Figure 4G) as well as the binding to the *PERK* promoter (Figure 4H). Together, these data demonstrate that CDC5L binds to the *PERK* promoter through its DBDs and represses *PERK* mRNA transcription in the nucleus.

### CDC5L activates the transcription of cell cycle genes but represses that of stress response genes

To further explore the function of CDC5L in regulating mRNA transcription, we carried out an RNA-sequencing (RNA-seq) analysis of the expression profile of HeLa cells treated with scrambled siRNA or si-*CDC5L.* We identified 1,226 differentially expressed genes (DEGs) that were downregulated and 871 DEGs that were upregulated in HeLa cells with *CDC5L* KD (fold change > 2 or < 0.5 and *p*-value < 0.05 in three repeats) (Figure 5A and Table S1). Functional analysis of the downregulated DEGs showed that the gene ontology (GO) terms were enriched in the regulation of cell cycle processes (Figure 5B), consistent with the essential role of CDC5L in cell cycle control as previously reported (Bernstein and Coughlin, 1998; Mu et al., 2014). Intriguingly, none of the upregulated DEGs in *CDC5L* KD cells were enriched in the regulation of the cell cycle; instead, the top-ranked GO terms of the upregulated DEGs were linked to stress response and the associated functions, such as “Response to ER stress”, “ER-nucleus signaling pathway”, and “Cellular response to stress” (Figure 5C).

**Figure 5.**
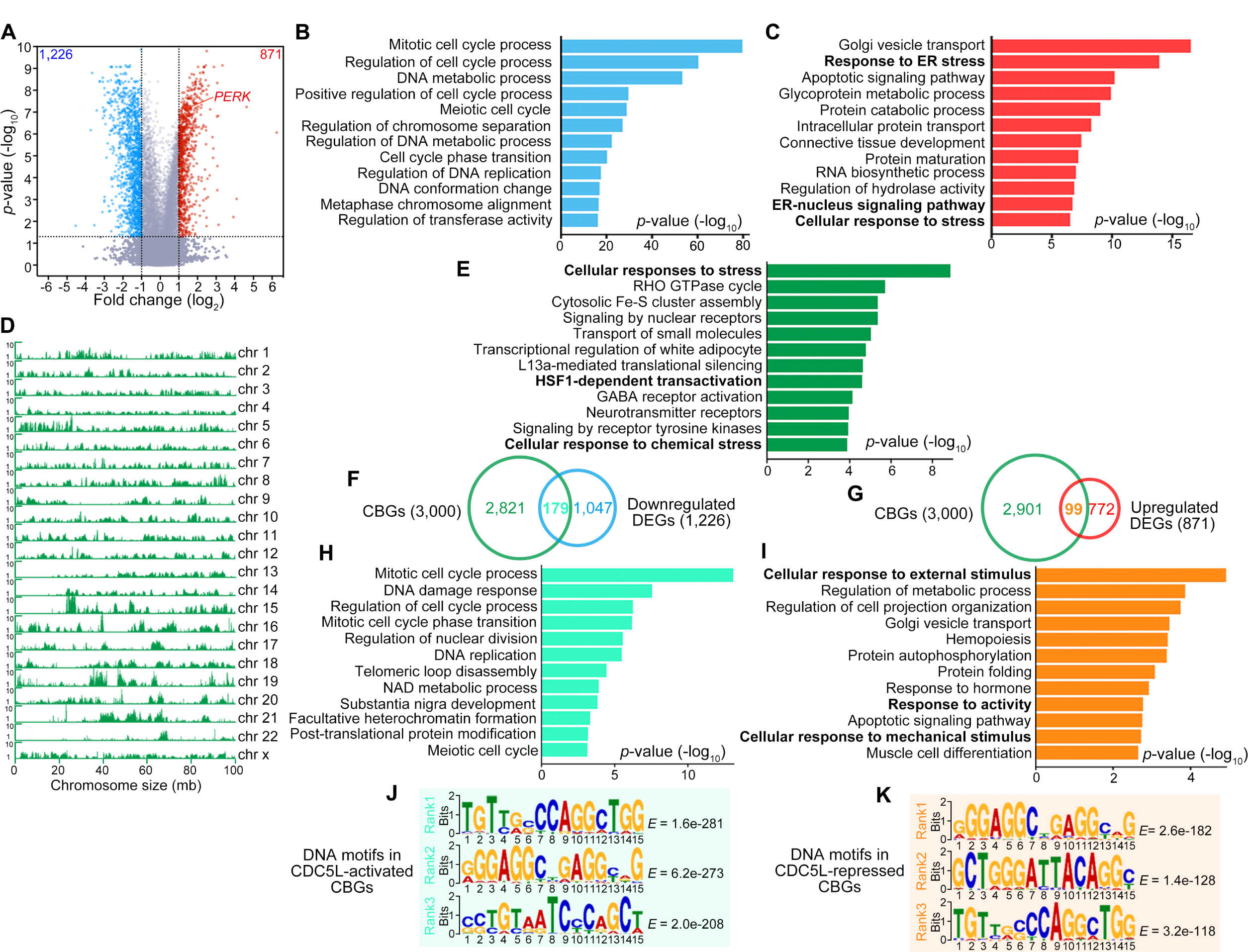
CDC5L activates the transcription of cell cycle genes but represses that of stress response genes. (**A**) Volcano plot showing the differentially expressed genes (DEGs) in the RNA-seq analysis of HeLa cells with *CDC5L* KD compared to the control cells (scrambled siRNA, si-Ctrl). Blue dots, downregulated DEGs; red dots, upregulated DEGs (fold change > 2 or < 0.5 and *p*-value < 0.05). (**B-C**) The top twelve enriched GO terms of biological processes associated with the downregulated (B) or upregulated (C) DEGs in (A). (**D**) The CDC5L-binding peaks throughout the genome of HeLa cells detected in the ChIP-seq analysis. (**E**) The top twelve enriched GO terms of the CDC5L-bound genes (CBGs) identified in the ChIP-seq. (**F-I**) Overlapping of the CBGs from the ChIP-seq (green) and the downregulated (F, blue) or upregulated (G, red) DEGs from the RNA-seq identifies 179 CDC5L-activated (cyan) and 99 CDC5L-repressed (orange) CBGs, and the top twelve enriched GO terms of these genes are shown in (H) and (I), respectively. (**J-K**) The top three-ranked DNA motifs for CDC5L binding in the CDC5L-activated (J) or CDC5L-repressed (K) CBGs.

As the DEGs identified in the RNA-seq analysis included genes both directly and indirectly regulated by *CDC5L*, we performed the ChIP sequencing (ChIP-seq) analysis of CDC5L in HeLa cells to investigate the genome-wide transcriptional targets directly bound by CDC5L (Figure 5D). 88,421 CDC5L-specific binding peaks were detected by MACS2 with a cutoff *q*-value < 0.05. The top 3,000 genes associated with the CDC5L-specific binding peaks were referred to as CDC5L-bound genes (CBGs) (Table S2). Functional analysis of the 3,000 CBGs identified “cellular responses to stress” as the most enriched biological process using Reactome (see Methods), along with “HSF1-dependent transactivation” and “cellular response to chemical stress” among the top twelve enriched pathways (Figure 5E).

We then cross-analyzed the data from the RNA-seq and ChIP-seq for overlapping genes to identify the direct targets of CDC5L. 179 CBGs in the ChIP-seq were found downregulated by *CDC5L* KD in the RNA-seq, which were the target genes transcriptionally activated by CDC5L (Tables S3 and Figures 5F); whereas 99 CBGs in the ChIP-seq were upregulated by *CDC5L* KD in the RNA-seq, which were the target genes transcriptionally repressed by CDC5L (Tables S4 and Figures 5G). The CDC5L-activated CBGs were enriched in cell cycle-related processes (Figure 5H), whereas CDC5L-repressed CBGs exhibited a strong association with pathways related to the regulation of stress response, including “Cellular response to external stimulus”, “Response to activity”, and “Cellular response to mechanical stimulus” (Figure 5I). And, the qPCR analysis confirmed the upregulation of other CDC5L-repressed stress response genes, such as *CALR, HEY1* and *BMP6*, in addition to *EIF2AK3* (*PERK*) in cells following *CDC5L* KD (Figure S8). Together, these results demonstrate that CDC5L acts as a transcription factor with dual function – it activates the transcription of cell cycle-related genes but represses the transcription of stress response genes.

Next, we analyzed the DNA motifs for CDC5L binding in the CDC5L-activated and CDCL-repressed CBGs using MEME-ChIP (see Methods). Notably, two of the top three DNA motifs, “GGGAGGCYGAGGCRG” (Y for T or C, R for G or A) and “TGTTGSCCAGGCTGG” (S for G or C), were shared in the CDC5L-activated and CDC5L-repressed CBGs (Figure 5J-K), suggesting that the opposite effects of CDC5L in regulating the transcription of cell cycle genes and stress response genes were not attained by binding to different DNA motifs. Alternatively, it might be achieved by CDC5L recruiting different transcription coactivators/corepressors in a lineage/stimulus-specific manner, which is worth further investigation in the future.

### KD of *CDC5L*/*Cdc5* reduces the stress tolerance in cell and fly models

As CDC5L acted as a transcriptional repressor of stress repronse genes, *CDC5L* KD led to upregulation of multiple stress signaling pathways, including the PERK-eIF2α pathway (as shown earlier in this study). It raised the question whether changes in stress sensitivity and response levels resulting from manipulation of *CDC5L* expression levels would confer a survival advantage or disadvantage. We then addressed this question using a propidium iodide (PI) staining assay, and the results showed that *CDC5L* KD enhanced whereas *CDC5L* OE ameliorated cell death in response to prolonged, mild stress (100 μM arsenite, 10 h) (Figure 6A-D). Importantly, changes in *CDC5L* expression levels did not cause significant cell death in the absence of stress, indicating that the effect was specific to stress rather than related to the function of CDC5L in the cell cycle or other processes.

**Figure 6.**
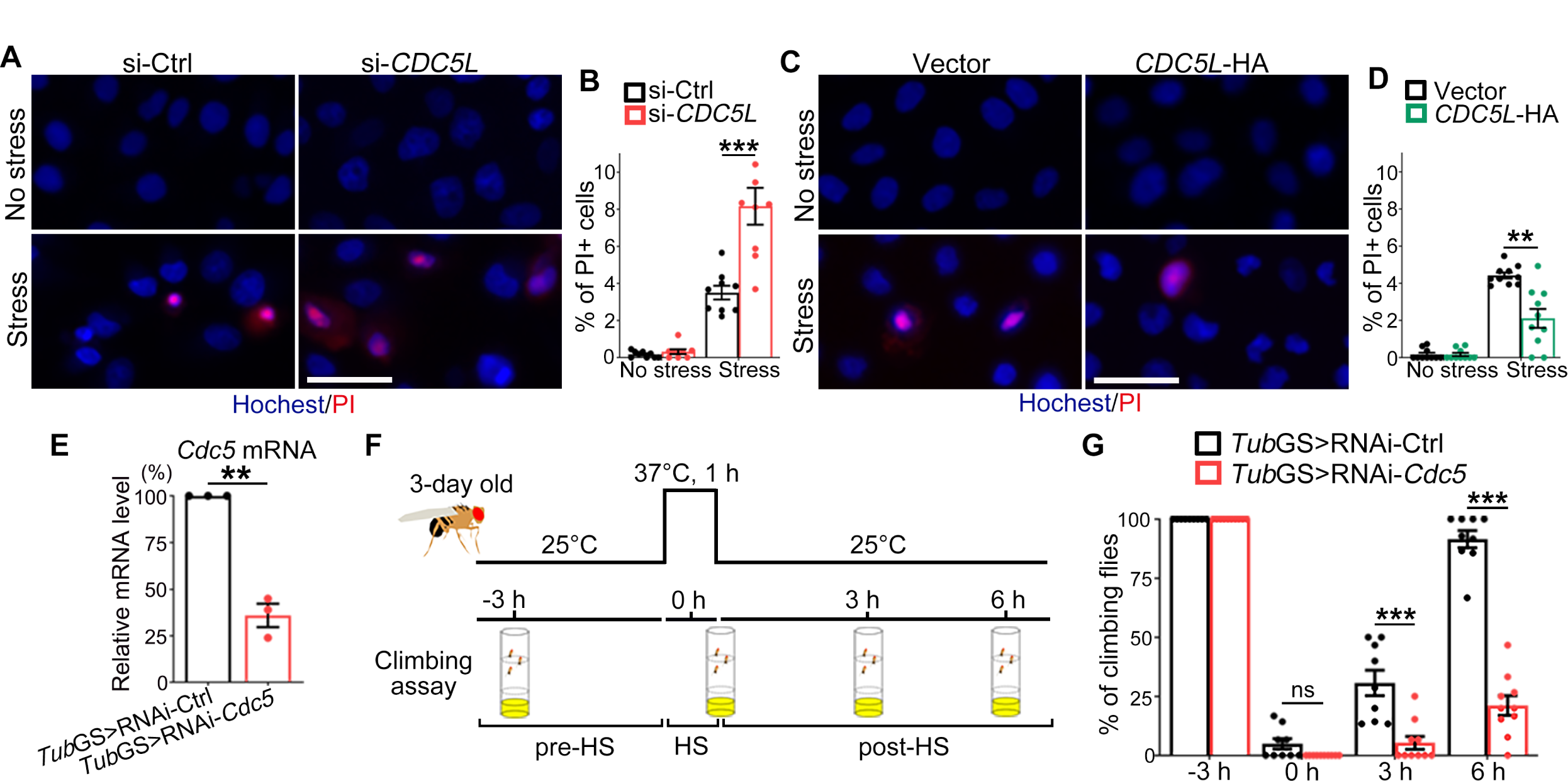
KD of *CDC5L*/*Cdc5* makes mammalian cells and flies less tolerant to mild stress. (**A**-**D**) Representative confocal images (A, C) and quantifications (B, D) of propidium iodide (PI) staining of HeLa cells with *CDC5L* KD (A-B) or OE (C-D) in response to prolonged, mild stress (100 μM arsenite, 10 h). Hoechst, nucleus; PI, cell death. (**E**) The qPCR analysis confirming the KD efficiency of RNAi-*Cdc5* in the *Tub*GS>RNAi-*Cdc5* flies. RNAi-Ctrl: RNAi-*mCherry* flies. The mRNA levels are normalized to *actin* and shown as the percentage to that of the *Tub*GS>RNAi-Ctrl flies. (**F**) A schematic diagram of the transient heat shock (HS) assay. 3-day old adult flies in tubes with food were gently submerged in a water bath of 37°C for 1 h and then put back to 25°C for recovery. The locomotor activity is assessed at 3 h before HS (pre-HS), right after HS (0 h), and at 3 or 6 h after HS (post-HS). (**G**) Quantification of the percentage of the *Tub*GS>RNAi-*Cdc5* flies in each vial that climb over 1 cm within 20 seconds at the indicated time points. Mean ± SEM; n ≥ 600 cells in each group from pooled results of 3 independent repeats in (B and D), n = 3 in (E), and n = 9-10 vials/group with about 15 flies per vial in (G). Student’s *t*-test; \*\**p* < 0.01, \*\*\**p* < 0.001; ns, not significant. Scale bars: 25 μm.

Finally, we extended our investigation to the *in vivo* model of *Drosophila*, the fruit fly. To avoid the influence on development, we used an inducible, ubiquitously expressed “*Tubulin* (*Tub*)-GeneSwitch (GS)” driver to downregulate *Cdc5* (the *Drosophila* homologue of *CDC5L*) in all fly cells from Day 1 of adulthood by adding RU486 to the fly food (Figure 6E). The flies, which were raised otherwise normally at 25°C, were subjected to a transient heat shock at 37°C for 1 h on Day 3 and then returned to 25°C for recovery (Figure 6F). To evaluate the viability of the flies, locomotor activity was measured using a climbing assay (see Methods). Before the heat shock (−3 h), the *Tub*GS>RNAi-*Cdc5* flies exhibited normal climbing capability similar to that of the *Tub*GS>RNAi-Ctrl flies (Figue 6G), which was consistent with the results that KD of *CDC5L* did not cause marked cell death without stress (Figure 6A-B). Right after the heat shock (0 h), both groups of the RNAi flies were unable to climb, confirming equal and sufficient heat-shock stress. At 3 h after the heat shock, the *Tub*GS>RNAi-Ctrl flies started to recover, with approximately 30% of thems able to climb, but the *Tub*GS>RNAi-*Cdc5* flies failed to climb; by 6 h after the heat shock, approximately 90% of the *Tub*GS>RNAi-Ctrl flies were able to climb, while only about 20% of the *Tub*GS>RNAi-*Cdc5* flies could climb (Figure 6G). Together, the *Tub*GS>RNAi-*Cdc5* flies appeared normal in the absence of stress; however, fewer of them recovered from heat shock and their recovery was significantly slower than that of the control flies, indicating that decrease in *Cdc5* expression levels made the flies less resistant to stress.

## DISCUSSION

In this study, we investigate the role of *CDC5L* in cellular stress and SGs, and demonstrate that CDC5L negatively regulates the stress signaling pathway PERK–eIF2α and prevents spontaneous and excessive activation of SG assembly. Formation of SGs is regarded as a common cellular mechanism to combat stress, and is conserved from yeast to flies and mammals (Protter and Parker, 2016; Zhang et al., 2018a; Grousl et al., 2022). Cellular toxins such as arsenite and physical conditions such as heat-shock stress are frequently used to induce SG assembly in laboratory research (including in part of this study), and there has been a coherent picture emerging for the signaling pathways and molecular mechanims that sense these stressors and activate SG assembly (Protter and Parker, 2016; Hofmann et al., 2021; Glauninger et al., 2022). Meanwhile, cells have evolved a plethora of mechanisms to eliminate cellular wastes and stressors in order to maintain homeostasis, and defects or LOF of a gene critical for maintaining the cell homeostasis may cause accumulation of cellular stress and activation of the stress response pathways (Berchtold et al., 2018; Corbet et al., 2021; Qifti et al., 2021).

Here, we report another layer of cellular mechanism regulating the stress response programs. We observe the occurrence of spontaneous SG formation upon downregulation of *CDC5L*, which is independent of its known functions in the cell cycle or the PRP19 spliceosome complex. Previous global proteomic analyses have provided indications of potential interactions between CDC5L and TDP-43, along with other ALS-associated RBPs (Freibaum et al., 2010; Llères et al., 2010; Youn et al., 2018). Nevertheless, our results show that CDC5L is predominantly localized to the nucleus and its protein levels are unaltered during stress. Nor is CDC5L translocated to the cytoplasm or recruited to SGs like TDP-43, FUS or TIAR. Hence, unlike the previous speculation, the CDC5L protein is not directly involved in the assembly of SGs and the interactions between CDC5L and the ALS-associated, SG-composing RBPs likely take place in the nucleus. Although our study does not exclude the possiblity that CDC5L may respond to stress at a post-translational level, the ChIP-PCR and luciferase reporter assays indicate that CDC5L binds to the *PERK* promoter and represses *PERK* transcription, which suppresses the PERK–eIF2α pathway and prevents spontaneous and excessive SG assembly.

Further investigation combining the RNA-seq and ChIP-seq analyses not only confirms the role of CDC5L as a transcription factor but also sheds light on its dual function in regulating two distinct gene clusters. One cluster of genes are predominantly linked to the regulation of the cell cycle and cell division. These genes display a significant downregulation in cells with *CDC5L* KD, emphasizing the crucial role of CDC5L in positively regulating the cell cycle (Bernstein and Coughlin, 1998; Mu et al., 2014). The other cluster comprises genes involved in cellular stress responses, such as *PERK*, which are upregulated in cells with *CDC5L* KD, indicating that CDC5L functions as a negative regulator of the stress response pathways. Adding to the intrigue, these distinct clusters of genes share the same top-ranked CDC5L-binding motifs within their promoters, suggesting that the differential transcriptional outcomes may be achieved through CDC5L’s recruitment of different transcription cofactors. Notably, the DBDs of CDC5L alone were sufficient and its nuclear localization was required for the repression of *PERK* transcription, suggesting that the nuclear co-repressors recruited by CDC5L might interact with CDC5L via 107-165 aa. It is possible that CDC5L interacts with different transcription cofactors and additional proteins to regulate the expression of different sets of genes involved in the cell cycle control and cellular stress responses. By utilizing these diverse transcriptional interactions, CDC5L may fine-tune gene expression and facilitate specific cellular functions in response to varying physiological conditions.

The primary functions of SGs are to regulate mRNA metabolism, protein synthesis and cellular stress responses. By sequestering and storing untranslated mRNAs as well as potentially harmful proteins, such as those activating apoptosis and pyroptosis (Tsai and Wei, 2010; Samir et al., 2019), SGs serve as a protective mechanism to prevent the translation of non-essential mRNAs and suppress the initation of cell death pathways during stress conditions (Anderson and Kedersha, 2009; Buchan and Parker, 2009; Protter and Parker, 2016; Alberti and Hyman, 2021; Glauninger et al., 2022). Intriguingly, upregulating stress response genes and promoting SG assembly by KD of *CDC5L* do not promote cell survival under stressed conditions. Instead, in both mammalian cells and the fly model, KD of *CDC5L*/*Cdc5* significantly reduces the ability of cells and individuals to withstand mild but prolonged stress, including stress induced by arsenite and heat shock. Thus, prompt and appropriate responses to cellular stress are crucial, and augmentation of stress signaling pathways or elevation of stress response levels do not necessarily provide enhanced cell adaptation or resilience to “prepare for” stress.

CDC5L has been found to be highly expressed in certain cancers and is believed to promote tumorigenesis by increasing cancer cell proliferation (Chen et al., 2016; Li et al., 2017; Zhang et al., 2020), in accordance with its known function in the cell cycle and cell division. In this study, we identify CDC5L as a transcriptional repressor that negatively regulates the PERK-eIF2α pathway and is required to prevent spontaneous SG assembly. Our study not only provides the first actual link between CDC5L and cellular stress but also presents a new layer of regulation and a gatekeeping mechanism for how cells surveil stress responses. The potential implications of alterations in CDC5L expression levels and defects in the *CDC5L* gene in disease pathogenesis are captivating and warrant further investigation in future studies.

## MATERIALS AND METHODS

### Plasmids and constructs

To generate the pCAG-*CDC5L-*HA and pCAG-*PERK-*HA plasmids, the DNA fragments encoding human CDC5L and PERK were amplified from cDNA of HeLa cells by RT-PCR using the specific primers (Table S5), and then inserted into a pCAG vector (a kind gift from Dr. Y. Chen) between the XhoI and EcoRI sites using the ClonExpress MultiS One Step Cloning Kit (Vazyme, C113). For construction of the pCAG-*CDC5L-ΔDBDs*-HA, pCAG-*CDC5L-ΔSAD*-HA, pCAG-*CDC5L-SAD*-HA, pCAG-*CDC5L-DBDs-*HA and pCAG-*CDC5L-DBDs-NES*-HA plasmids, the truncated CDC5L fragments were amplified from the above pCAG-*CDC5L-*HA plasmid by PCR with the designated primers (Table S5) and sub-cloned into the pCAG vector, as described above.

To generate the pCAG-*GFP-G3BP1* plasmid, the DNA fragment encoding human G3BP1 was amplified from cDNA of HeLa cells by RT-PCR and then inserted into the pCAG-*GFP-TDP-43* plasmid (Wang et al., 2020) using the aforementioned method.

To generate the pcDNA3.1-*PERK-luc* plasmid, the promoter region (from −2 kb to 189 bp of the transcription start site) of *PERK* was amplified from the genomic DNA of HeLa cells by PCR and then sub-cloned into the pcDNA3.1-*luciferase* (firefly) plasmid (a kind gift from Dr. K. He) between the MluI and NheI sites to replace original promoter using the ClonExpress MultiS One Step Cloning Kit (Vazyme, C113). All constructs were confirmed by sequencing before use. The sequences of all the primers used for plasmid construction are summarized in Supplemental Table S5.

### Cell culture and transfection

HeLa cells (American Type Culture Collection, CCL-2) were cultured in the Dulbecco’s modified Eagle’s medium (DMEM) (Basalmedia, L110KJ) supplemented with 10% (vol/vol) fetal bovine serum (FBS) (Sigma-Aldrich, F8318) and 1% penicillin/streptomycin at 37°C in a humidified atmosphere of 95% air and 5% CO2. Transient transfection of siRNA oligonucleotides (Genepharma, Shanghai) or plasmids was performed using Lipofectamine RNAiMAX (Thermo Fisher, 13778150) in Opti-MEM (Thermo Fisher, 51985034) or the PolyJet In Vitro DNA Transfection Reagent (SignaGen Laboratories, SL100688) in DMEM according to the manufacturers’ instructions. Cells were transfected for at least 48 h before subsequent pharmacological treatment or examination.

The siRNA oligonucleotides used in this study are listed below:

si-Ctrl (scrambled siRNA): 5’-UUCUCCGAACGUGUCACGUTT-3’

si-*hCDC5L*#1: 5’-GCUGGAAGAACGUGAAAUATT-3’

si-*hCDC5L*#2: 5’-GCUCUCAAGUGAAGCUUAUTT-3’

si-*hPRP19*: 5’-GCCAAGUUCAUCGCUUCAATT-3’

si-*hPLRG1*: 5’-GCUGCAGAACCACAAAUUATT-3’

si-*hBCAS2*: 5’-GCUCGACAACCAAUUGAAUTT-3’

si-*hPERK*: 5’-GUGGAAAGGUGAGGUAUAUTT-3’

si-*hG3BP1*: 5’-CCUGAUGAUUCUGGAACUUTT-3’

si-*hG3BP2*: 5’-CAGUGAAUGUCAUACUAAATT-3’

### Immunocytochemistry and confocal imaging

HeLa cells grown on coverslips in a 24-well plate were transfected and/or treated as indicated, and then sequentially fixed with 4% paraformaldehyde (Sangon, A500684-0500) in the phosphate-buffered saline (PBS) (30 min), permeabilized in 0.5% Triton X-100 (Sigma-Aldrich, T8787) in the PBS (PBST; 30 min), and blocked with 3% goat serum in the PBST (blocking buffer; 60 min) at room temperature (RT). Thereafter, cells were probed with the specific primary and secondary antibodies in the blocking buffer at 4°C overnight. After washing three times in the PBST, the cells were then mounted in the VECTASHIELD Antifade Mounting Medium with DAPI (Vector Laboratories, H-1200) on glass slides. Confocal images were taken with the Leica TCS SP8 confocal microscopy system using a 63x oil objective (NA = 1.4) and analyzed with Leica Application Suite X (LAS X) software. The images were processed and assembled into figures using Adobe Photoshop 2021.

### Cellular stress assays

Arsenite stress: HeLa cells were transfected with the indicated siRNA or plasmids for at least 48 h before treated with 250 μM of NaAsO2 or PBS for the indicated time periods up to 30 min, followed by fixation with 4% paraformaldehyde. For the recovery experiment, culture medium containing NaAsO2 was removed and the cells were incubated in fresh medium for the indicated time prior to fixation.

Heat-shock stress: HeLa cells were transfected with the indicated siRNA or plasmids for at least 48 h before transferred to an incubator of 42°C for the indicated durations of time up to 30 min, followed by fixation. For the recovery experiment, the cell plates were returned to the regular cell incubator of 37°C, and the cells were fixed and examined at the indicated time points.

### Fluorescence recovery after photobleaching (FRAP) assay

The FRAP assay was performed using the FRAP module of the Leica SP8 confocal microscopy system. In brief, each GFP-G3BP1 granule was bleached using a 488 nm laser at 100% laser power for approximately 4 s. After photobleaching, time-lapse images were captured every 1.2 s for the about 1 min. For each indicated time point (t), the fluorescence intensity within the bleached granule was normalized to the fluorescence intensity of a nearby, unbleached granule (to control for photobleaching during live imaging). The normalized fluorescence intensity of pre-bleaching was set to 100% and the normalized fluorescence intensity at each time point (It) was used to calculate the fluorescence recovery (FR) according to the following formula: FR(t) = It/Ipre-bleaching. GraphPad Prism was used to plot and analyze the FRAP experiments.

### Protein extraction and western blotting

Total proteins were extracted from cells using a 2% SDS extraction buffer (50mM Tris pH 7.4, 2% SDS, 3% DTT, 40% glycerol and 0.04% bromophenol blue) containing the protease inhibitor cocktail (Roche, 04693132001) and the phosphatase inhibitor cocktail (Roche, 04906837001). For separation of soluble and insoluble proteins, cells were lysed on ice in a RIPA buffer (50 mM Tris pH 8.0, 150 mM NaCl, 1% NP-40, 5 mM EDTA, 0.5% sodium deoxycholate, 0.1% SDS) supplemented with protease and phosphatase inhibitors. After sonication, the homogenates were centrifuged at 13,000 *g* for 20 min at 4°C. The supernatant was used as the soluble fraction and the pellets containing the insoluble fraction were dissolved in a urea buffer (9 M urea, 50 mM Tris buffer, pH 8.0) after washing.

All protein samples were then boiled at 100°C for 10 min, separated using 8% Tris-glycine sodium dodecyl-sulfate-polyacrylamide gel electrophoresis (SDS-PAGE), and probed with the primary and secondary antibodies listed below. The immunoblots were detected using the High-sig ECL Western Blotting Substrate (Tanon, 180-5001). Images were captured using an Amersham Imager 600 (GE Healthcare) and densitometry was measured using Image Quant TL Software (GE Healthcare). The contrast and brightness were optimized equally using Adobe Photoshop CS2021. GAPDH was used as a loading control for normalization, as indicated in the figures.

### Antibodies

The following primary antibodies were used in this study: rabbit anti-PERK (CST, 3192S), rabbit anti-eIF2α (CST, 5324S), rabbit anti-p-eIF2α (CST, 9721S), rabbit anti-HA (CST, 3724T), rabbit anti-TIAR (CST, 8509S), mouse anti-CDC5L (BD Biosciences, 612362), mouse anti-G3BP1 (BD Biosciences, 611127), mouse anti-G3BP1 (Proteintech, 66486-1), mouse anti-GAPDH (Proteintech, 60004-1), normal rabbit IgG (CST, 2729S). HRP conjugated secondary antibodies: goat anti-mouse (Sigma-Aldrich, A4416) and goat anti-rabbit (Sigma-Aldrich, A9169). Fluorescent secondary antibodies: goat anti mouse-Alexa Fluor 568 (Thermo Fisher, A11031) and goat anti rabbit-Alexa Fluor 488 (Thermo Fisher, A11034).

### Cell cycle synchronization

Cell cycle synchronization was performed as previously described (Hong et al., 2004; Thuy et al., 2017). In brief, log-phase cells were first incubated with 1.7 μM 20-hydroxyecdysone (Selleck, S2417) for 24 h to obtain cells in the G2 phase. Cells were then rinsed three times with the PBS, resuspended in fresh DMEM supplemented with 10% FBS along with 1.5 mM hydroxyurea (Selleck, S1896), and cultured for 18 h to reach the G1/S phase. Afterward, these cells were rinsed three times with the PBS, cultured in the above medium without hydroxyurea, and harvested at the indicated time points for the subsequent flow cytometry analysis or confocal imaging. For the flow cytometry analysis (Moflo Astrioseq EQ, Beckman), the cells were fixed in ethanol, incubated with RNase A and treated with propidium iodide (PI) (Sangon, E607306) for 30 min.

### RNA extraction and quantitative real-time PCR (qPCR)

For qPCR analysis, total RNA was isolated from HeLa cells or flies using RNAiso Plus (Takara, 9109) according to the manufacturer’s instruction. After DNase (Yeasen, 11141-B) treatment to eliminate genomic DNA, the reverse transcription (RT) reactions were performed using Hifair® III 1st Strand cDNA Synthesis SuperMix for qPCR (Yeasen, 11141ES60). The resulting cDNA was then used for real-time qPCR with the Taq Pro Universal SYBR qPCR Master Mix (Vazyme, Q712-02) using the Quant Studio™ 6 Flex Real-Time PCR system (Thermo Fisher). The mRNA levels of *GAPDH* or *actin* were used as an internal control to normalize the mRNA levels of genes of interest. The qPCR primers used in this study are summarized in Table S5.

### Dual-luciferase reporter assay

The *PERK-luc* (firefly) and pcDNA3.1-*rluc* (*Renilla*) plasmids were co-transfected into HeLa cells along with the indicated siRNA or plasmids in 96-well plates. After transfection for 48 h, the cells were examined using the Dual-Glo® Luciferase Assay System (Promega, E2920) according to the manufacturer’s instruction. The luminescence signals of the firefly and *Renilla* luciferases were measured using the BioTek Synergy2 Multi-Detection Microplate Reader.

### Chromatin immunoprecipitation (ChIP) and ChIP-PCR

HeLa cells (about 6 x 10^7^ cells/6 cm dish) transfected with the indicated plasmids were fixed in 1% (vol/vol) formaldehyde/PBS at 37°C for 10 min and quenched with 125 mM glycine at RT for 5 min. After washing, the samples were lysed in a cytoskeleton buffer (10 mM PIPES (pH 6.8), 100 mM NaCl, 3 mM MgCl2, 1 mM EGTA (pH 7.6), 0.3 M sucrose, 0.5% Triton X-100, 0.5 mM DTT and 5 mM sodium butyrate) at 4°C for 10 min, followed by centrifugation at 400 *g* at 4°C for 5 min. The pellets were resuspended in a micrococcal nuclease (MNase) buffer (50 mM Tris-HCl (pH 7.5), 4 mM MgCl2, 1 mM CaCl2, 0.3 M sucrose, 0.5 mM DTT and 5 mM sodium butyrate) with 2,000 units MNase (Beyotime, D7201S) at 37°C for 20 min, and the digestion was then stopped with 0.5 M EDTA at 4°C for 2 min.

The digested chromatin samples were centrifuged at 13,000 *g* at 4°C for 5 min and resuspended in 150 μL of a ChIP lysis buffer (1% SDS, 50 mM Tris-HCl, pH 8.0 and 10 mM EDTA) at 4°C for 10 min. 1,350 μL of the RIPA buffer was added to each lysed samples and sonicated for 3 cycles (10 s on/10 s off) using the Medical Ultrasonic Homogenizer Processor Cell Disruptor Mixer (Jingxin, JY92-IIN), resulting in DNA fragments of 100-600 bp. The supernatants were used for immunoprecipitation with Protein G Dynabeads (Thermo Fisher, 10004D) associated with the 0.35 μg anti-HA rabbit mAb (CST, 3724T) or normal rabbit IgG (CST, 2729S) at 4°C for 5 h with gentle rotation.

Immunoprecipitated DNAs were dissolved in the ChIP Elution Buffer (0.1% SDS, 50 mM EDTA, 50 mM Tris-HCl, pH 8.0 and 50 mM NaHCO3) and reverse cross-linked at 65°C overnight. The DNAs were then purified with the QIAGEN QIAquick PCR Purification Kit (QIAGEN, 28106) and amplified using the *PERK* or *GAPDH* promoter primers (Table S5). The resulting DNA products (195 bp for *PERK* and 166 bp for *GAPDH*) were examined by electrophoresis on 1.5% agarose gels.

### RNA-seq and data analysis

Total RNAs from HeLa cells (about 5 x 10^7^ cells/10 cm dish) transfected with scrambled siRNA (si-Ctrl) or si-*CDC5L* were extracted using RNAiso Plus (Takara, 9109) according to the manufacturer’s instruction. The quality and quantity of the RNA samples were examined using the 5400 Fragment Analyzer (Agilent, M5312AA). RNA libraries were constructed utilizing the NEBNext® Ultra RNA Library Prep Kit for Illumina (NEB, E7530L) and sequenced on an Illumina NovaSeq 6000 PE150 platform using the 150-bp pair-end sequencing parameters (Novogene, Beijing).

For the RNA-seq data, the output gene count tables from Salmon v0.9.1 (Patro et al., 2017) based on alignments to the human genome (hg38) annotation were used as input into the limma package v3.16.2 (Law et al., 2014). The differential expression genes (DEGs) were identified with the cutoff as follows: p-value < 0.05 and fold change > 2 or < 0.5. To analyze Gene Ontology (GO) and pathway enrichment for select subsets of genes, Metascape (https://metascape.org/) was used and *p*-value < 0.01 was considered significant.

### ChIP-seq and data analysis

The ChIP samples were prepared as described above. For sequencing, the quality and quantity of the immunoprecipitated DNAs were assessed using the Qubit®Fluorometer (Thermo Fisher, Q32866) and agarose gel electrophoresis. The ChIP libraries were constructed utilizing NEBNext® Ultra™ II DNA Library Prep Kit (NEB, E7645L) and sequenced on an Illumina NovaSeq 6000 PE150 platform using the 150-bp pair-end sequencing parameters (Novogene, Beijing).

For the ChIP-seq data, paired-end reads were filtered for redundant reads and aligned using Bowtie2 (v2.2.5) to the human genome (hg38). Broad peaks were called with the MACS2 v2.2.7.1 software using input as a negative control with a cutoff of *q*-value (minimum false discovery rate) < 0.05. 88,421 peaks were called by MACS2, followed by blacklist filtering. The peaks identified by MACS2 were then assigned to the associated genes using BETA minus (http://cistrome.org/BETA/index.html) for downstream analyses. The top 3,000 assigned CDC5L-bound genes (CBGs) were uploaded to the online platform Metascape (https://metascape.org/) to identify the Reactome pathways with a cutoff of *p*-value < 0.01. The CDC5L-bound genomic regions were uploaded to MEME-ChIP v5.5.4 (Machanick and Bailey, 2011) for motif analysis with a cutoff of *E*-value (adjusted *p*-value) < 0.05.

### Cell death assessment by PI staining

HeLa cells were seeded at a density of 1×10^5^ cells/well in 24-well plates and transfected with the indicated siRNAs or plasmids for 48 h. To detect dead cells, the PI Staining Kit (Sangon, E607306) was used at 1:1,000 dilution and incubated at 37°C for 60 min according to the manufacturer’s instruction.

### *Drosophila* strains and husbandry

The RNAi-*Cdc5* (TH03978.N) fly strain was obtained from the TsingHua Fly Center (THFC), the RNAi-*mCherry* (#35785, a control line for short hairpin RNAi KD) obtained from the Bloomington Drosophila Stock Center (BDSC), and the *Tub*GS was a kind gift from N. Bonini. All flies were raised on standard cornmeal media and maintained at 25°C and 60% relative humidity. To induce the expression of the *Tub*GS driver, adult flies were raised on regular fly food supplemented with 160 μg/mL RU486 (mifepristone; TCI, 84371-65-3) dissolved in ethanol.

### Transient heat shock assay in flies

About 15 flies in each regular fly vial with cornmeal food at the bottom were submerged into a water bath of 37°C for 1 h and then placed back to the fly incubator at 25°C for recovery. The locomotor activity, indicative of fly viability, was measured through a climbing assay at the indicated time points before or after the heat shock. In brief, flies were transferred into an empty polystyrene vial and gently tapped down to the bottom. The number of flies that climbed over a distance of 1 cm within 20 seconds was recorded. The test was repeated three times for each vial and 9-10 vials were examined per group.

### Statistical analysis

Statistical significance in this study is determined by one-way ANOVA with Tukey’s HSD post-hoc test, or unpaired, two-tailed Student’s *t*-test at \**p* < 0.05, \*\**p* < 0.01 and \*\*\**p* < 0.001. Error bars represent the standard error of the mean (SEM).

## ACKNOWLEDGEMENTS

We thank the *Drosophila* stock centers BDSC and THFC providing the fly strains, Drs A. Li, Y. Chen, K. He and N. Bonini for sharing reagents, plasmids and fly lines, members of the Fang lab for helpful discussion, and A. Li and C. Tang for comments and critical reading of the manuscript. This study was supported by grants from the National Natural Science Foundation of China (32325016, 32270812, 22337005 and 31970697) and the Science and Technology Commission of Shanghai Municipality (20490712600 and 2019SHZDZX02) to Y. Fang.

## AUTHOR CONTRIBUTIONS

Y.D. and Y.F. conceived the research; B.Q., S.L., Y.D., and Y.F. designed the experiments; B.Q., S.L., Y.D., J.C. and Q.W. performed the experiments; R.H. and W.Y. contributed important new reagents; B.Q., S.L., F.Q., W.L. and Y.F. analyzed the data and interpret the results; B.Q. and S.L. prepared the figures; and B.Q., S.L., and Y.F. wrote the paper. All authors read and approved the final manuscript.

## AVAILABILITY OF DATA AND MATERIAL

All essential data are presented in the main manuscript and the online Supporting Information. The RNA-seq and ChIP-seq datasets are deposited in the GEO Datasets (accession# 242280) and will be made publicly available when the paper is published. All unique and stable reagents generated in this study are available from the corresponding author with a complete Material Transfer Agreement.

## SUPPLEMENTAL FIGURES

**Figure S1.**
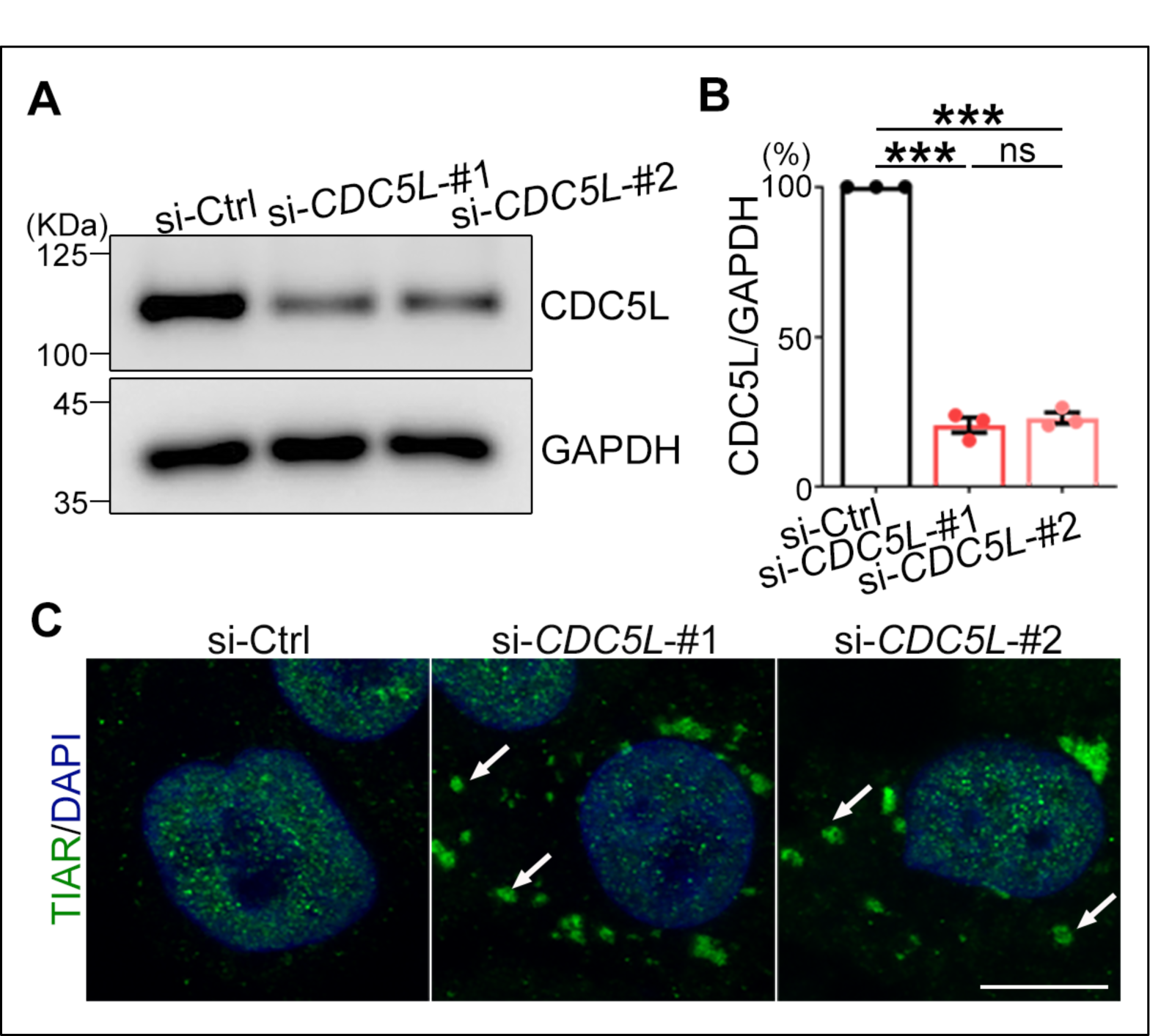
KD of *CDC5L* induces spontaneous SG formation. (**A**) Western blot analysis confirms the decrease in CDC5L protein levels in HeLa cells using the two independent siRNAs targeting *CDC5L*. (**B**) Quantification of relative CDC5L protein levels (normalized to GAPDH) in (A). si-Ctrl, scrambled control siRNA. (**C**) Immunostaining for another SG marker, TIAR, confirms the spontaneous formation of SGs (indicated by arrows) in HeLa cells with *CDC5L* KD. DAPI, nucleus. Mean ± SEM, n = 3. One-way ANOVA; \*\*\**p* < 0.001; ns, not significant. Scale bar: 10 μm.

**Figure S2.**
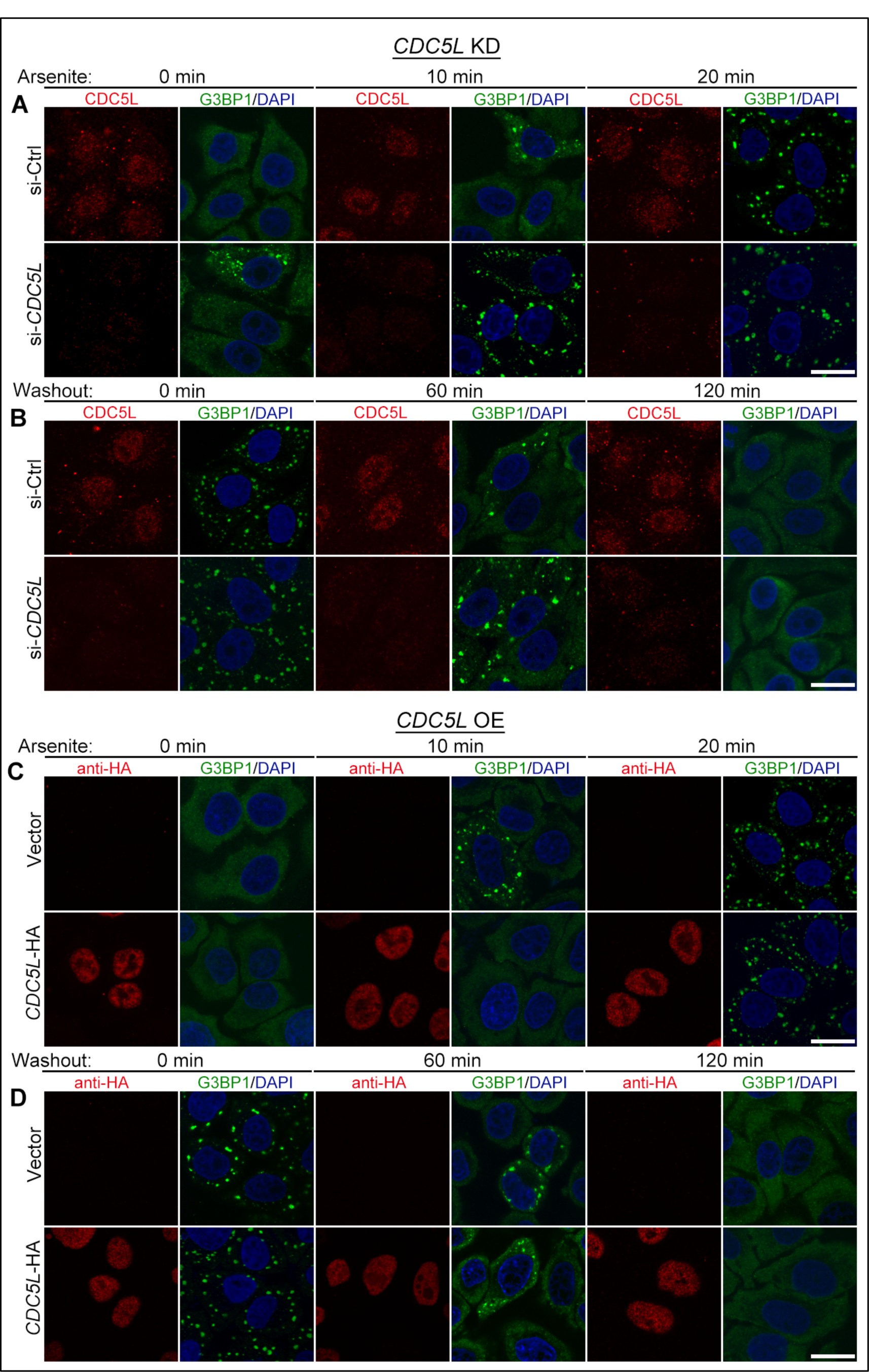
*CDC5L* levels impact the assembly and disassembly of SGs induced by arsenite stress. (**A-D**) Representative confocal images at the indicated time points over the course of assembly (A, C) and disassembly (B, D) of arsenite-induced SGs in HeLa cells following siRNA KD of *CDC5L* (A-B), or transient OE of *CDC5L*-HA (C-D). Quantifications are presented in Figure 1I, K, M, and O. DAPI, nucleus. Scale bars: 20 μm.

**Figure S3.**
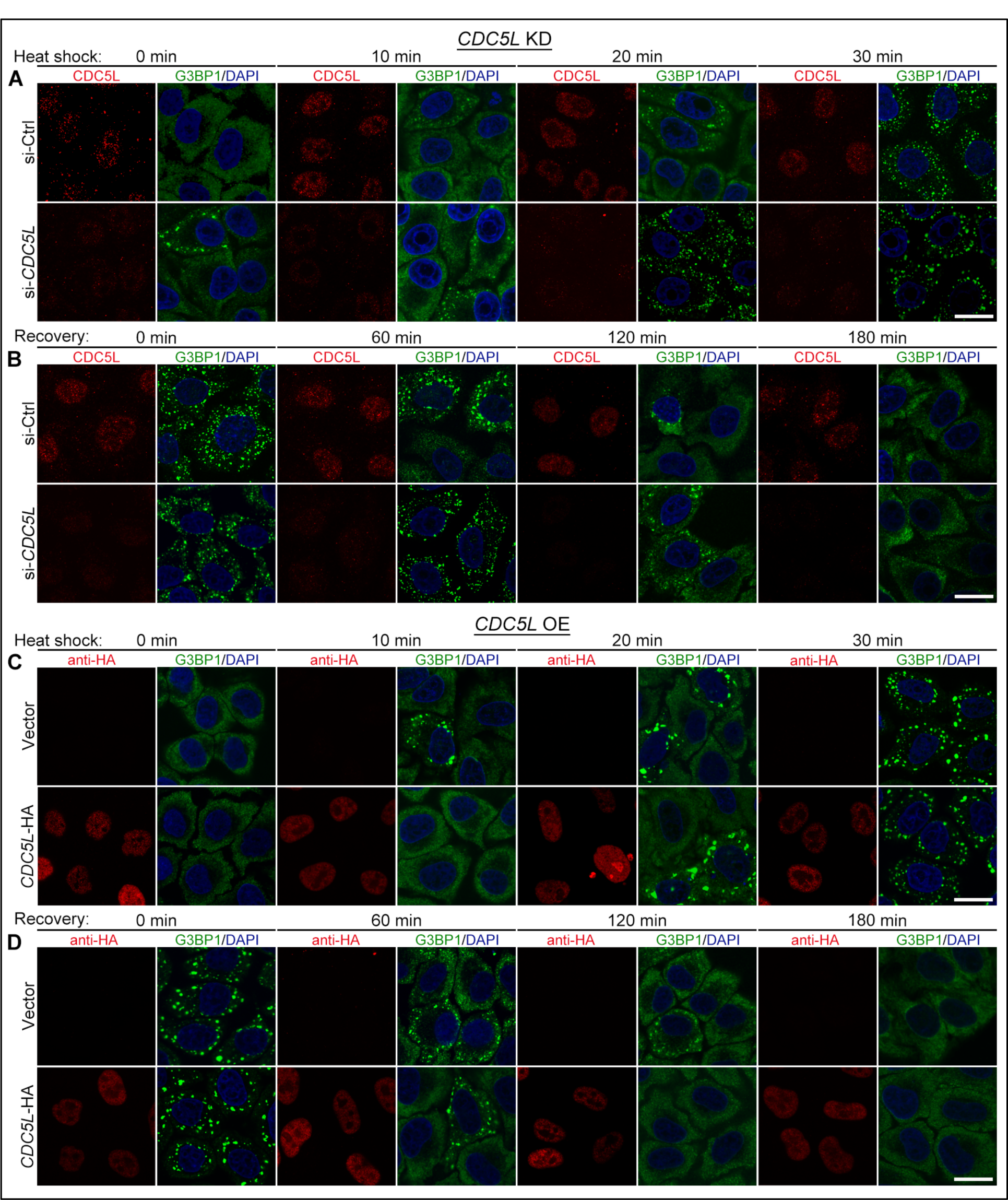
*CDC5L* levels impact the assembly and disassembly of SGs induced by heat-shock stress. (**A-D**) Representative confocal images at the indicated time points over the course of assembly (A, C) and disassembly (B, D) of heat shock-induced SGs in HeLa cells following siRNA KD of *CDC5L* (A-B), or transient OE of *CDC5L*-HA (C-D). Quantifications are presented in Figure 1J, L, N, and P. DAPI, nucleus. Heat shock: 42°C; recovery: 37°C. Scale bars: 20 μm.

**Figure S4.**
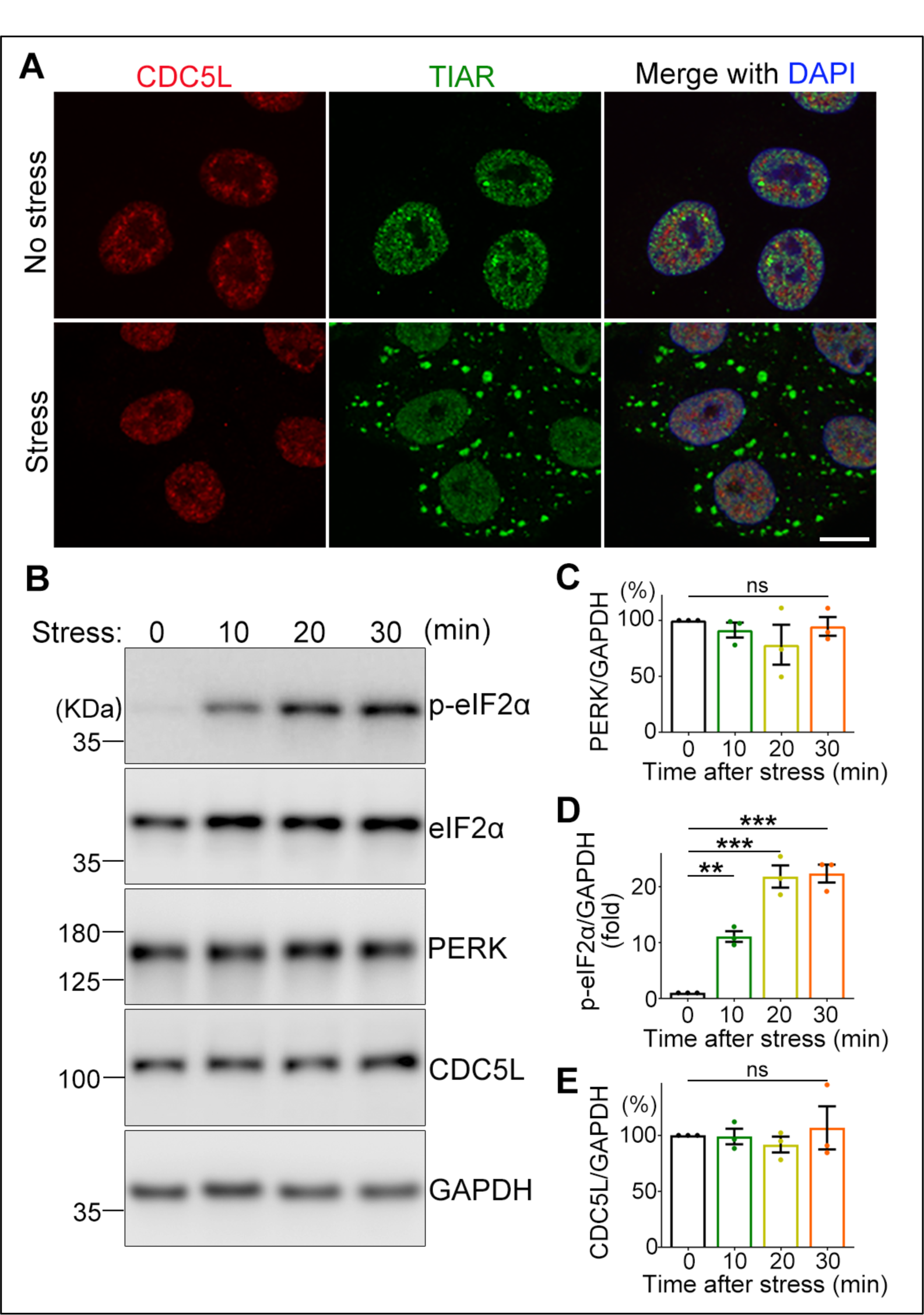
CDC5L does not translocate to SGs in the cytoplasm or increase its expression levels in response to stress. (**A**) Representative confocal images of HeLa cells untreated or treated with arsenite (250 μM, 30 min). Cells are stained for CDC5L, TIAR (SGs), and DAPI (nucleus). (**B-E**) Western blot analysis (B) and quantifications (C-E) of the relative protein levels of PERK (C), phosphorylated eIF2α (p-eIF2α) (D), and CDC5L (E) at the indicated time points during arsenite stress (250 μM). Protein levels are normalized to GAPDH and shown as percentage relative to the “no stress” group (0 min; set to 100%). Mean ± SEM, n = 3. One-way ANOVA; \*\**p* < 0.01, \*\*\**p* < 0.001; ns, not significant. Scale bar: 10 μm.

**Figure S5.**
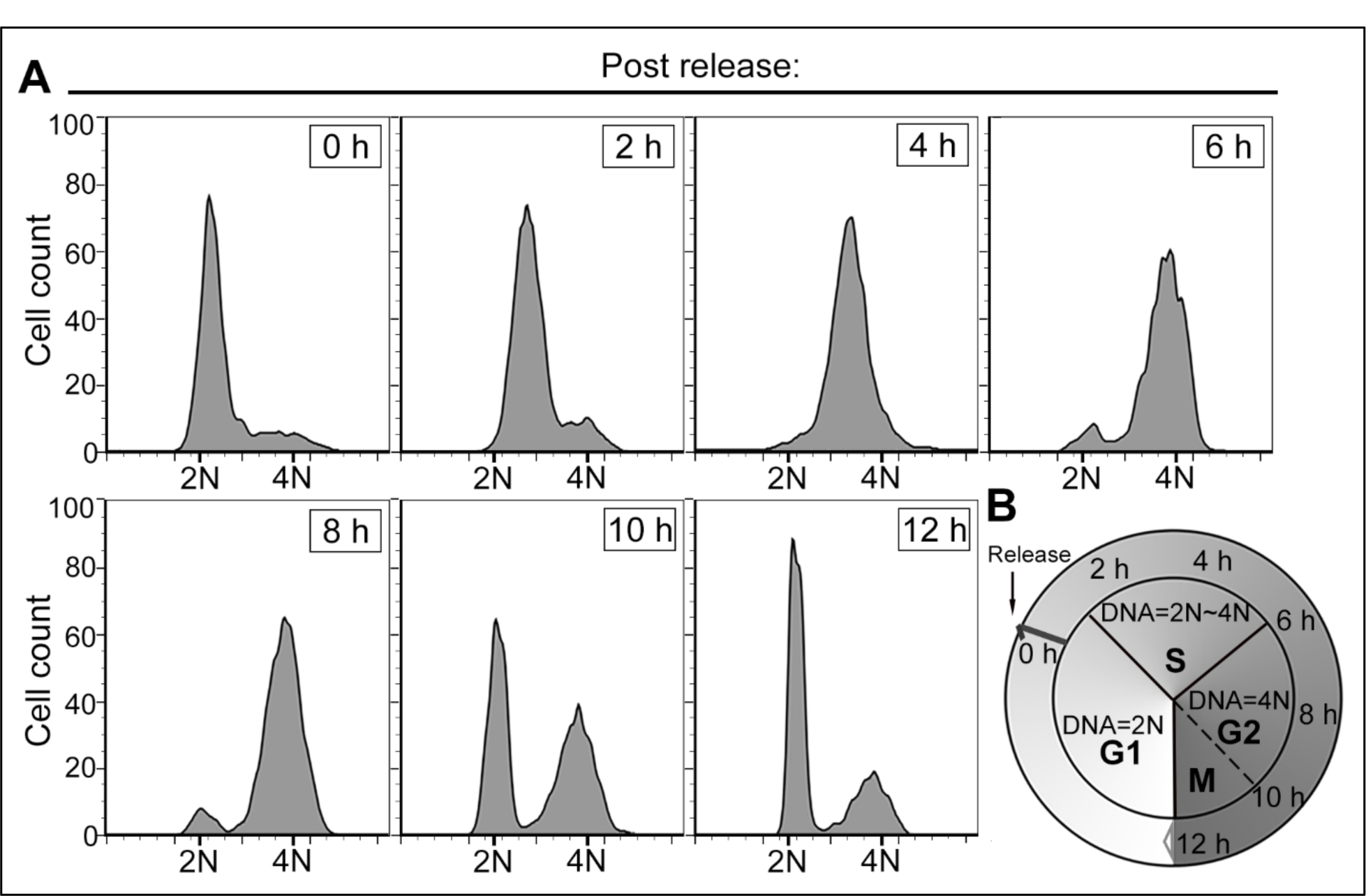
Determining the different phases in the cell cycle assay. (**A**) The different phases of the cell cycle are determined by flow cytometry analysis. Briefly, HeLa cells are synchronized and arrested at the G1 phase (DNA content = 2N) using hydroxyurea. Upon hydroxyurea removal from the culture medium, cells are released from the G1 and transition to the S phase (DNA content = 2N∼4N) within 2-6 h, progress to the G2/M phase (DNA content = 4N) within 6-10 h, and then enter a new cycle starting with the G1 phase (DNA content = 2N) by approximately 12 h. (**B**) A schematic of the cell cycle assay illustrating the different phases and their corresponding time points as determined in (A). G1 phase: the growth and metabolism phase; S phase: the DNA replication phase; G2 phase: the growth of structural elements for the mitosis phase; M phase, the mitosis phase.

**Figure S6.**
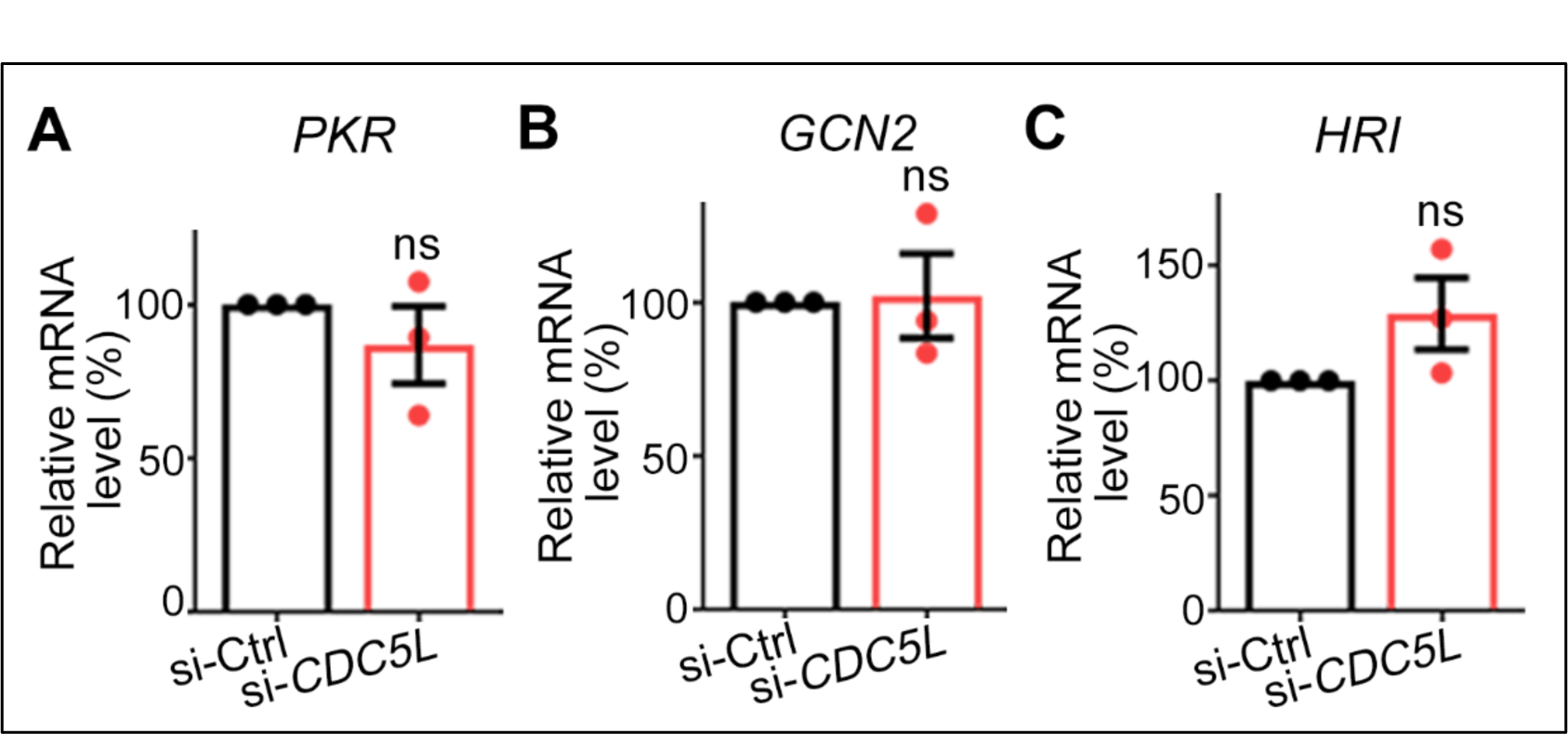
KD of *CDC5L* does not affect the mRNA levels of the other three eIF2α kinases. (**A-C**) qPCR analysis of the mRNA levels of the other three eIF2α kinases—*PKR* (A), *GCN2* (B), and *HRI* (C)—in HeLa cells with *CDC5L* KD. mRNA levels are normalized to *GAPDH* and shown as percentage to the si-Ctrl group (scrambled siRNA). Mean ± SEM, n = 3. Student’s *t*-test; ns, not significant.

**Figure S7.**
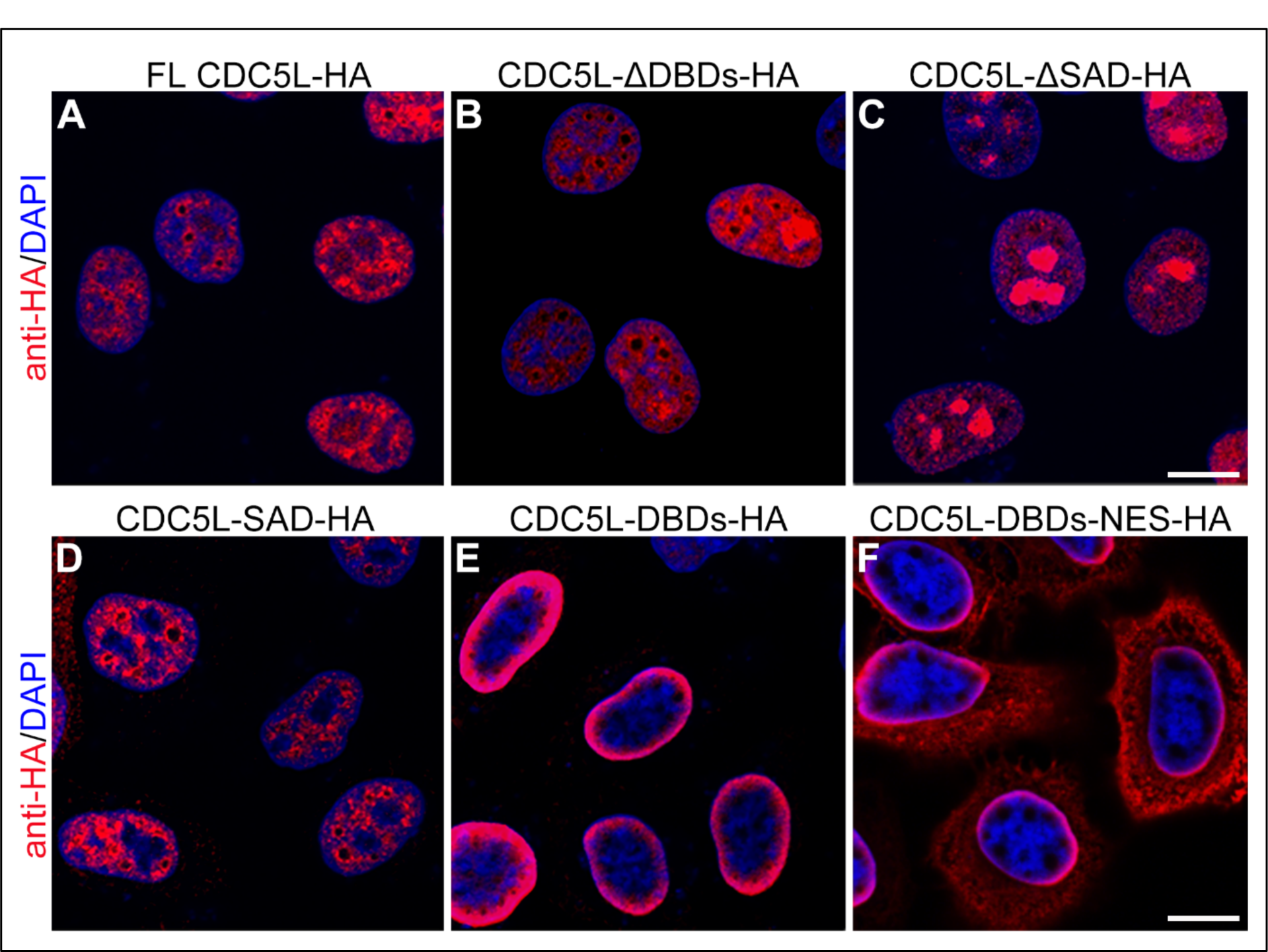
Expression of full-length (FL) and truncated CDC5L proteins in HeLa cells. Representative confocal images depicting the subcellular distribution of FL (A) and truncated CDC5L-HA proteins in HeLa cells, including ΔDBDs (B), ΔSAD (C), SAD (D), DBDs (E) and DBDs-NES (F). Cells are stained for HA and DAPI (nucleus). Scale bars: 10 μm. Of note, although CDC5L-DBDs-HA lacks the putative NLS and shows a different sub-nuclear distribution from the other CDC5L truncation proteins, it is still localized within the nucleus, mostly along the inner side of the nuclear membrane, as demarcated by DAPI staining (E). The addition of an NES to the DBDs results in the cytoplasm localization (F), which abolishes the binding of DBDs to the *PERK* promoter as well as the repression on *PERK* transcription by DBDs, as demonstrated in Figure 4G-4H.

**Figure S8.**
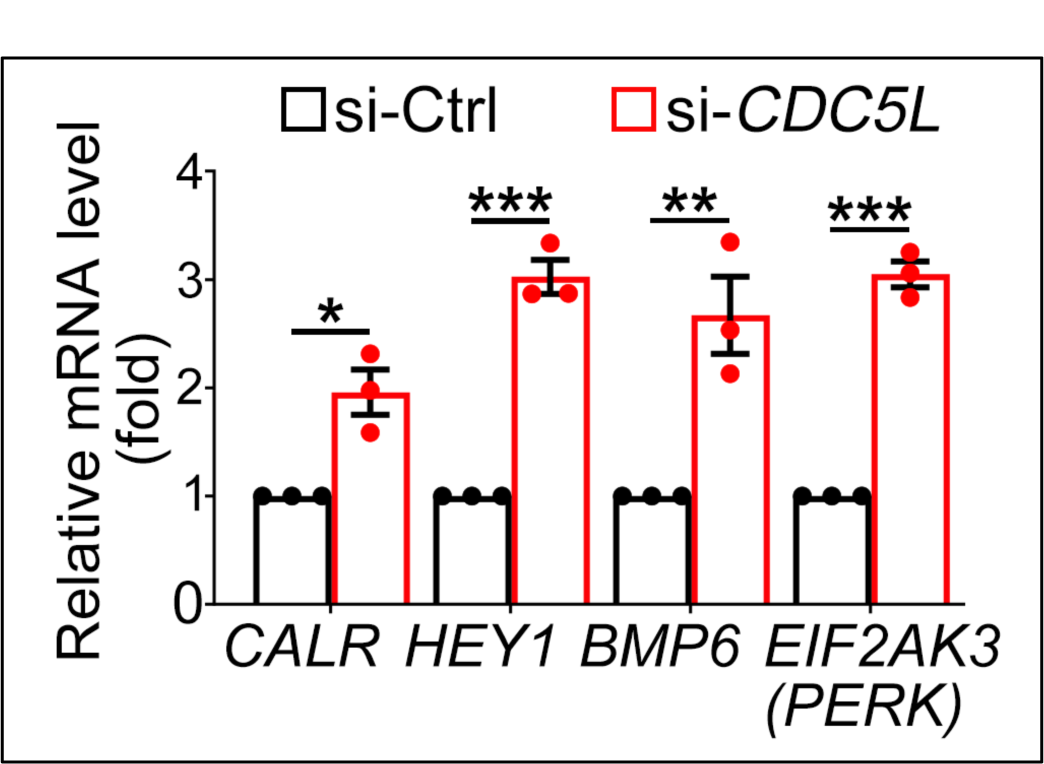
Confirmation of the upregulation of other CDC5-repressed, stress response genes in cells following *CDC5L* KD. The results of the qPCR analysis confirms the upregulation of other CDC5L-repressed, direct target genes involved in stress responses, such as *CALR*, *HEY1* and *BMP6*, in addition to *EIF2AK3* (*PERK*) in HeLa cells following *CDC5L* KD. All mRNA levels are normalized to *GAPDH* and shown as fold change to that of the control group (set to 1). si-Ctrl: scrambled siRNA. Mean ± SEM, n = 3. Student’s *t*-test; \**p* < 0.05, \*\**p* < 0.01, \*\*\**p* < 0.001.

## SUPPLEMENTAL TABLES

**Table S1. The DEGs in RNA-seq analysis of HeLa cells with CDC5L KD**

**Table S2. The CBGs from ChIP-seq analysis of HeLa cell expressing CDC5L-HA**

**Table S3. The downregulated CBGs and their GO term analysis**

**Table S4. The upregulated CBGs and their GO term analysis**

**Table S5. Primer information for PCR and qPCR in this study**

## REFERENCE

Ajuh, P., Kuster, B., Panov, K., Zomerdijk, J.C., Mann, M., and Lamond, A.I. (2000). Functional analysis of the human CDC5L complex and identification of its components by mass spectrometry. EMBO J 19, 6569–6581.

Alberti, S., Gladfelter, A., and Mittag, T. (2019). Considerations and challenges in studying liquid-liquid phase separation and biomolecular condensates. Cell 176, 419–434.

Alberti, S., and Hyman, A.A. (2021). Biomolecular condensates at the nexus of cellular stress, protein aggregation disease and ageing. Nat Rev Mol Cell Biol 22, 196–213.

Anderson, P., and Kedersha, N. (2009). Stress granules. Curr Biol 19, R397–398.

Bernstein, H.S., and Coughlin, S.R. (1998). A mammalian homolog of fission yeast Cdc5 regulates G2 progression and mitotic entry. J Biol Chem 273, 4666–4671.

Baradaran-Heravi, Y., Van Broeckhoven, C., and van der Zee, J. (2020). Stress granule mediated protein aggregation and underlying gene defects in the FTD-ALS spectrum. Neurobiology of Disease 134, 104639.

Berchtold, D., Battich, N., and Pelkmans, L. (2018). A systems-level study reveals regulators of membrane-less organelles in human cells. Mol Cell 72, 1035–1049.e1035.

Boudrez, A., Beullens, M., Groenen, P., Van Eynde, A., Vulsteke, V., Jagiello, I., Murray, M., Krainer, A.R., Stalmans, W., and Bollen, M. (2000). NIPP1-mediated interaction of protein phosphatase-1 with CDC5L, a regulator of pre-mRNA splicing and mitotic entry. J Biol Chem 275, 25411–25417.

Buchan, J.R., and Parker, R. (2009). Eukaryotic stress granules: the ins and outs of translation. Mol Cell 36, 932–941.

Chanarat, S., and Strasser, K. (2013). Splicing and beyond: the many faces of the Prp19 complex. Biochim Biophys Acta 1833, 2126–2134.

Chen, W., Zhang, L., Wang, Y., Sun, J., Wang, D., Fan, S., Ban, N., Zhu, J., Ji, B., and Wang, Y. (2016). Expression of CDC5L is associated with tumor progression in gliomas. Tumour Biol 37, 4093–4103.

Corbet, G.A., Burke, J.M., and Parker, R. (2021). ADAR1 limits stress granule formation through both translation-dependent and translation-independent mechanisms. J Cell Sci 134, jcs258783.

Donnelly, N., Gorman, A.M., Gupta, S., and Samali, A. (2013). The eIF2alpha kinases: their structures and functions. Cell Mol Life Sci 70, 3493–3511.

Fang, M.Y., Markmiller, S., Vu, A.Q., Javaherian, A., Dowdle, W.E., Jolivet, P., Bushway, P.J., Castello, N.A., Baral, A., Chan, M.Y., Linsley, J.W., Linsley, D., Mercola, M., Finkbeiner, S., Lecuyer, E., Lewcock, J.W., and Yeo, G.W. (2019). Small-molecule modulation of TDP-43 recruitment to stress granules prevents persistent TDP-43 accumulation in ALS/FTD. Neuron 103, 802–819.e11.

Freibaum, B.D., Chitta, R.K., High, A.A., and Taylor, J.P. (2010). Global analysis of TDP-43 interacting proteins reveals strong association with RNA splicing and translation machinery. Journal of Proteome Research 9, 1104–1120.

Glauninger, H., Wong Hickernell, C.J., Bard, J.A.M., and Drummond, D.A. (2022). Stressful steps: Progress and challenges in understanding stress-induced mRNA condensation and accumulation in stress granules. Mol Cell 82, 2544–2556.

Grousl, T., Vojtova, J., Hasek, J., and Vomastek, T. (2022). Yeast stress granules at a glance. Yeast 39, 247–261.

Guillen-Boixet, J., Kopach, A., Holehouse, A.S., Wittmann, S., Jahnel, M., Schlussler, R., Kim, K., Trussina, I., Wang, J., Mateju, D., Poser, I., Maharana, S., Ruer-Gruss, M., Richter, D., Zhang, X., Chang, Y.T., Guck, J., Honigmann, A., Mahamid, J., Hyman, A.A., Pappu, R.V., Alberti, S., and Franzmann, T.M. (2020). RNA-Induced conformational switching and clustering of G3BP drive stress granule assembly by condensation. Cell 181, 346–361.e17.

Gwon, Y., Maxwell, B.A., Kolaitis, R.-M., Zhang, P., Kim, H.J., and Taylor, J.P. (2021). Ubiquitination of G3BP1 mediates stress granule disassembly in a context-specific manner. Science 372, eabf6548.

Hans, F., Glasebach, H., and Kahle, P.J. (2020). Multiple distinct pathways lead to hyperubiquitylated insoluble TDP-43 protein independent of its translocation into stress granules. J Biol Chem 295, 673–689.

Hofmann, S., Kedersha, N., Anderson, P., and Ivanov, P. (2021). Molecular mechanisms of stress granule assembly and disassembly. Biochim Biophys Acta Mol Cell Res 1868, 118876.

Hong, S.H., Hong, B., Kim, D.C., Rho, M.S., Park, J.I., Rha, S.H., Jun, H.S., and Jeong, J.S. (2004). Involvement of mitogen-activated protein kinases and p21Waf1 in hydroxyurea-induced G1 arrest and senescence of McA-RH7777 rat hepatoma cell line. Exp Mol Med 36, 493–498.

Kumar, R., and Haider, S. (2022). Protein network analysis to prioritize key genes in amyotrophic lateral sclerosis. IBRO Neurosci Rep 12, 25–44.

Law, C.W., Chen, Y., Shi, W., and Smyth, G.K. (2014). voom: Precision weights unlock linear model analysis tools for RNA-seq read counts. Genome Biol 15, R29.

Li, J., Zhang, N., Zhang, R., Sun, L., Yu, W., Guo, W., Gao, Y., Li, M., Liu, W., Liang, P., Deng, W., and Cui, X. (2017). CDC5L promotes hTERT expression and colorectal tumor growth. Cell Physiol Biochem 41, 2475–2488.

Llères, D., Denegri, M., Biggiogera, M., Ajuh, P., and Lamond, A.I. (2010). Direct interaction between hnRNP-M and CDC5L/PLRG1 proteins affects alternative splice site choice. EMBO Rep 11, 445–451.

Machanick, P., and Bailey, T.L. (2011). MEME-ChIP: motif analysis of large DNA datasets. Bioinformatics 27, 1696–1697.

Mu, R., Wang, Y.B., Wu, M., Yang, Y., Song, W., Li, T., Zhang, W.N., Tan, B., Li, A.L., Wang, N., et al. (2014). Depletion of pre-mRNA splicing factor Cdc5L inhibits mitotic progression and triggers mitotic catastrophe. Cell Death Dis 5, e1151.

Patro, R., Duggal, G., Love, M.I., Irizarry, R.A., and Kingsford, C. (2017). Salmon provides fast and bias-aware quantification of transcript expression. Nat Methods 14, 417–419.

Portz, B., Lee, B.L., and Shorter, J. (2021). FUS and TDP-43 phases in health and disease. Trends Biochem Sci 46, 550–563.

Protter, D.S.W., and Parker, R. (2016). Principles and properties of stress granules. Trends Cell Biol 26, 668–679.

Putnam, A., Thomas, L., and Seydoux, G. (2023). RNA granules: functional compartments or incidental condensates? Genes Dev 37, 354–376.

Qifti, A., Jackson, L., Singla, A., Garwain, O., and Scarlata, S. (2021). Stimulation of phospholipase Cbeta1 by Galphaq promotes the assembly of stress granule proteins. Sci Signal 14, eaav1012.

Ravanidis, S., Kattan, F.G., and Doxakis, E. (2018). Unraveling the pathways to neuronal homeostasis and disease: mechanistic insights into the role of RNA-binding proteins and associated factors. Int J Mol Sci 19, 2280.

Roden, C., and Gladfelter, A.S. (2021). RNA contributions to the form and function of biomolecular condensates. Nat Rev Mol Cell Biol 22, 183–195.

Samir, P., Kesavardhana, S., Patmore, D.M., Gingras, S., Malireddi, R.K.S., Karki, R., Guy, C.S., Briard, B., Place, D.E., Bhattacharya, A., Sharma, B.R., Nourse, A., King, S.V., Pitre, A., Burton, A.R., Pelletier, S., Gilbertson, R.J., and Kanneganti, T.D. (2019). DDX3X acts as a live-or-die checkpoint in stressed cells by regulating NLRP3 inflammasome. Nature 573, 590–594.

Sherf, B. A., Navarro, S. L., Hannah, R. R. & Wood, K. V. (1996). Dual-luciferase^TM^ reporter assay: an advanced co-reporter technology integrating firefly and renilla luciferase assays. Promega Notes 57, 2–9.

Sivan, G., Kedersha, N., and Elroy-Stein, O. (2007). Ribosomal slowdown mediates translational arrest during cellular division. Mol Cell Biol 27, 6639–6646.

Sun, H., Huang, Y., Mei, S., Xu, F., Liu, X., Zhao, F., Yin, L., Zhang, D., Wei, L., Wu, C., et al. (2021). A nuclear export signal is required for cGAS to sense cytosolic DNA. Cell Rep 34, 108586.

Thuy, T.T., Tam, N.T., Anh, N.T., Hau, D.V., Phong, D.T., Thang, L.Q., Adorisio, S., Van Sung, T., and Delfino, D.V. (2017). 20-Hydroxyecdysone from Dacrycarpus imbricatus bark inhibits the proliferation of acute myeloid leukemia cells. Asian Pac J Trop Med 10, 157–159.

Tsai, N.-P., and Wei, L.-N. (2010). RhoA/ROCK1 signaling regulates stress granule formation and apoptosis. Cellular Signalling 22, 668–675.

van Maldegem, F., Maslen, S., Johnson, C.M., Chandra, A., Ganesh, K., Skehel, M., and Rada, C. (2015). CTNNBL1 facilitates the association of CWC15 with CDC5L and is required to maintain the abundance of the Prp19 spliceosomal complex. Nucleic Acids Res 43, 7058–7069.

Wang, C., Duan, Y., Duan, G., Wang, Q., Zhang, K., Deng, X., Qian, B., Gu, J., Ma, Z., Zhang, S., Guo, L., Liu, C., and Fang, Y. (2020). Stress induces dynamic, cytotoxicity-antagonizing TDP-43 nuclear bodies via paraspeckle lncRNA NEAT1-mediated liquid-liquid phase separation. Mol Cell 79, 443–458 e447.

Williams, S.D., Zhu, H., Zhang, L., and Bernstein, H.S. (2006). Adenoviral delivery of human CDC5 promotes G2/M progression and cell division in neonatal ventricular cardiomyocytes. Gene Therapy 13, 837–843.

Wolozin, B., and Ivanov, P. (2019). Stress granules and neurodegeneration. Nat Rev Neurosci 20, 649–666.

Yang, P., Mathieu, C., Kolaitis, R.M., Zhang, P., Messing, J., Yurtsever, U., Yang, Z., Wu, J., Li, Y., Pan, Q., Yu, J., Martin, E.W., Mittag, T., Kim, H.J., and Taylor, J.P. (2020). G3BP1 is a tunable switch that triggers phase separation to assemble stress granules. Cell 181, 325–345.e28.

Youn, J.Y., Dunham, W.H., Hong, S.J., Knight, J.D.R., Bashkurov, M., Chen, G.I., Bagci, H., Rathod, B., MacLeod, G., Eng, S.W.M., Angers, S., Morris, Q., Fabian, M., Cote, J.F., and Gingras, A.C. (2018). High-density proximity mapping reveals the subcellular organization of mRNA-associated granules and bodies. Mol Cell 69, 517–532 e511.

Youn, J.Y., Dyakov, B.J.A., Zhang, J., Knight, J.D.R., Vernon, R.M., Forman-Kay, J.D., and Gingras, A.C. (2019). Properties of stress granule and P-body proteomes. Mol Cell 76, 286–294.

Zhang, K., Coyne, A.N., and Lloyd, T.E. (2018a). Drosophila models of amyotrophic lateral sclerosis with defects in RNA metabolism. Brain Research 1693, 109–120.

Zhang, K., Daigle, J.G., Cunningham, K.M., Coyne, A.N., Ruan, K., Grima, J.C., Bowen, K.E., Wadhwa, H., Yang, P., Rigo, F., et al. (2018b). Stress granule assembly disrupts nucleocytoplasmic transport. Cell 173, 958–971.e17.

Zhang, Y., Liu, T., Meyer, C.A., Eeckhoute, J., Johnson, D.S., Bernstein, B.E., Nusbaum, C., Myers, R.M., Brown, M., Li, W., et al. (2008). Model-based analysis of ChIP-Seq (MACS). Genome Biol 9, R137.

Zhang, Z., Mao, W., Wang, L., Liu, M., Zhang, W., Wu, Y., Zhang, J., Mao, S., Geng, J., and Yao, X. (2020). Depletion of CDC5L inhibits bladder cancer tumorigenesis. J Cancer 11, 353–363.

